# The 3D enhancer network of the developing T cell genome is controlled by SATB1

**DOI:** 10.1101/2021.07.09.451769

**Authors:** Tomas Zelenka, Antonios Klonizakis, Despina Tsoukatou, Sören Franzenburg, Petros Tzerpos, Dionysios-Alexandros Papamatheakis, Ioannis-Rafail Tzonevrakis, Christoforos Nikolaou, Dariusz Plewczynski, Charalampos Spilianakis

## Abstract

Mechanisms of tissue-specific gene expression regulation via spatial coordination of gene promoters and distal regulatory elements are still poorly understood. We investigated the 3D genome organization of developing murine T cells and identified SATB1, a tissue-specific genome organizer, enriched at the anchors of promoter-enhancer chromatin loops. We assessed the function of SATB1 in T cell chromatin organization and compared it to the conventional genome organizer CTCF. SATB1 builds a more refined layer of genome organization upon a CTCF scaffold. To understand the regulatory implications of SATB1 loopscape structure, we generated *Satb1*^fl/fl^*Cd4*-Cre^+^ (*Satb1* cKO) conditional knockout animals which suffered from autoimmunity. We aimed to identify molecular mechanisms responsible for the deregulation of the immune system in *Satb1* cKO animals. H3K27ac HiChIP and Hi-C experiments indicated that SATB1 primarily mediates promoter-enhancer loops affecting master regulator genes (such as *Bcl6*), the T cell receptor locus and adhesion molecule genes, collectively being critical for cell lineage specification and immune system homeostasis. Our findings unravel the function of a tissue-specific factor that controls transcription programs, via spatial chromatin arrangements complementary to the chromatin structure imposed by ubiquitously expressed genome organizers.

## Introduction

In order to store the large amount of genetic information, higher eukaryotes developed spatial and functional genome organization into compartments and domains. The A and B compartments represent the largest organizational units and they functionally correspond to active and inactive chromatin regions, respectively (Lieberman-Aiden et al., 2009; Rao et al., 2014). These compartments are further partitioned into topologically associated domains (TADs; Dixon et al., 2012; Nora et al., 2012), although due to their heterogeneous nature, new terminology better reflecting the reality is slowly being adopted (Rowley and Corces, 2018). Structural segmentation of chromatin in mammals is driven by architectural proteins such as CTCF and the cohesin complex (Nora et al., 2017; Rao et al., 2017; Schwarzer et al., 2017). Depletion of either CTCF (Nora et al., 2017), cohesin (Rao et al., 2017) or its loading factor *Nipbl* (Schwarzer et al., 2017) leads to global disruption of TAD organization, yet surprisingly with only modest transcriptional changes and unaffected A/B compartments. This, together with the recent findings at base-pair resolution (Hua et al., 2021), indicates the presence of additional mechanisms of the three-dimensional (3D) chromatin organization. The elimination of cohesin loading factor *Nipbl* unveiled a finer compartment structure that reflected the underlying epigenetic landscape (Schwarzer et al., 2017). This observation is in line with the model in which the primary driver of chromatin organization is the actual transcriptional state (Rowley et al., 2017). Indeed, RNA polymerase II and transcription itself are tightly linked to the formation of finer-scale structures of chromatin organization (Hsieh et al., 2020). Nonetheless, transcriptional inhibition has only a modest effect on promoter-enhancer contacts (Hsieh et al., 2020). Similarly, the inhibition of BET proteins, degradation of BRD4 or dissolution of transcriptional phase condensates all yield in the disrupted transcription, however they also have just a little impact on promoter-enhancer interactions (Crump et al., 2021). In contrast, an experimental disruption of TADs (Lupiáñez et al., 2015) or direct manipulation of promoter-enhancer contacts (Deng et al., 2012) both result in alterations of gene expression. Additionally, chromatin reorganization often precedes changes in transcription during development and differentiation (Apostolou et al., 2013; Stadhouders et al., 2018), suggesting that in many scenarios 3D genome organization instructs the transcriptional programs. However, the precise mechanisms on how chromatin organization is linked to gene expression regulation, especially in the context of cell lineage specification, still remain poorly understood. Several transcription factors have been shown to mediate promoter-enhancer interactions and thus also the underlying transcriptional programs in a tissue-specific manner and often even independent of CTCF and cohesin (Giammartino et al., 2020; Kim and Shendure, 2019; Stadhouders et al., 2019). Such an additional regulatory layer of chromatin organization, provided by transcription factors, may represent the missing link between high order chromatin structure and transcriptional regulation.

In this work, we aimed to identify drivers of regulatory chromatin loops in developing murine T cells, as a great model of tissue-specific gene expression regulation. We have identified SATB1, a factor exhibiting enriched occupancy at gene promoters and enhancers involved in long-range chromatin interactions. SATB1 has been attributed to many biological roles, mostly during T cell development (Zelenka and Spilianakis, 2020), but it also regulates the function of cell types such as the epidermis (Fessing et al., 2011) and neurons (Balamotis et al., 2012; Denaxa et al., 2012). Moreover, SATB1 is also overexpressed in a wide array of cancers and is positively associated with increased tumour size, metastasis, tumour progression, poor prognosis and reduced overall survival (Sunkara et al., 2018). Originally described as a Special AT-rich Binding protein (Dickinson et al., 1992), it is known for its propensity to bind DNA regions with more negative torsional stress (Ghosh et al., 2019). In a proposed model, SATB1 dimers bound to DNA interact with each other to form a tetramer in order to mediate long-range chromatin loops (Wang et al., 2012, 2014). The regulatory function of SATB1 is controlled by post-translational modifications (Kumar et al., 2006) and indirectly also by protein-protein interactions with chromatin modifying complexes (Fujii et al., 2003; Jangid et al., 2014; Kumar et al., 2005; Purbey et al., 2009; Yasui et al., 2002).

To understand the principles of chromatin organization in murine thymocytes and their impact on physiology, we performed Hi-C and HiChIP experiments and compared the roles of tissue-specific SATB1 and ubiquitously expressed CTCF genome organizers. Our findings were complemented by ATAC-seq, RNA-seq and H3K27ac HiChIP experiments in WT and *Satb1* cKO thymocytes to further unravel the functional roles of SATB1. This represents a comprehensive genome-wide study, systematically probing all SATB1-mediated chromatin loops in the T cell nucleus. A number of datasets combined with unbiased analytical approaches indicated the presence of a functional organizational layer built upon a general chromatin scaffold mediated by conventional genome organizers, such as CTCF, specifically regulating expression of master regulator genes and adhesion molecule genes essential for proper T cell development.

## Results

### Detection of regulatory chromatin loops in T cells

In order to unravel the active promoter-enhancer connectome in T cells, we performed H3K27ac HiChIP experiments (Mumbach et al., 2016) in C57BL/6J (WT) thymocytes. Loop calling at 5 kbp resolution (FDR ≤ 0.01) yielded 16,458 regulatory loops. To identify the prospective protein factors associated with these regulatory loops, we intersected the anchors of these loops with all the available murine ChIP-seq datasets from blood cells, using the enrichment analysis of ChIP-Atlas (Oki et al., 2018). The most highly enriched protein factors included RAG1, RAG2, BCL11b, SATB1 and TCF1 (Figure 1A). Both RAG1/2 proteins are known to be associated with the H3K27ac histone modification (Maman et al., 2016; Teng et al., 2015), however their main known role relies in the recombination of B and T cell receptor loci (Fugmann et al., 2000). BCL11b and TCF1 are well-studied factors specifying T cell lineage commitment, whose roles in forming the chromatin landscape in T cells have been recently addressed (Emmanuel et al., 2018; Garcia-Perez et al., 2020; Hu et al., 2018; Johnson et al., 2018). We drew our attention to SATB1, which displayed significant enrichment at the H3K27ac loop anchors (Figure 1B) and also represents a known genome organizer (Cai et al., 2003, 2006), yet with a limited number of genome-wide studies targeting its role in 3D chromatin organization of T cells.

**Figure 1.**
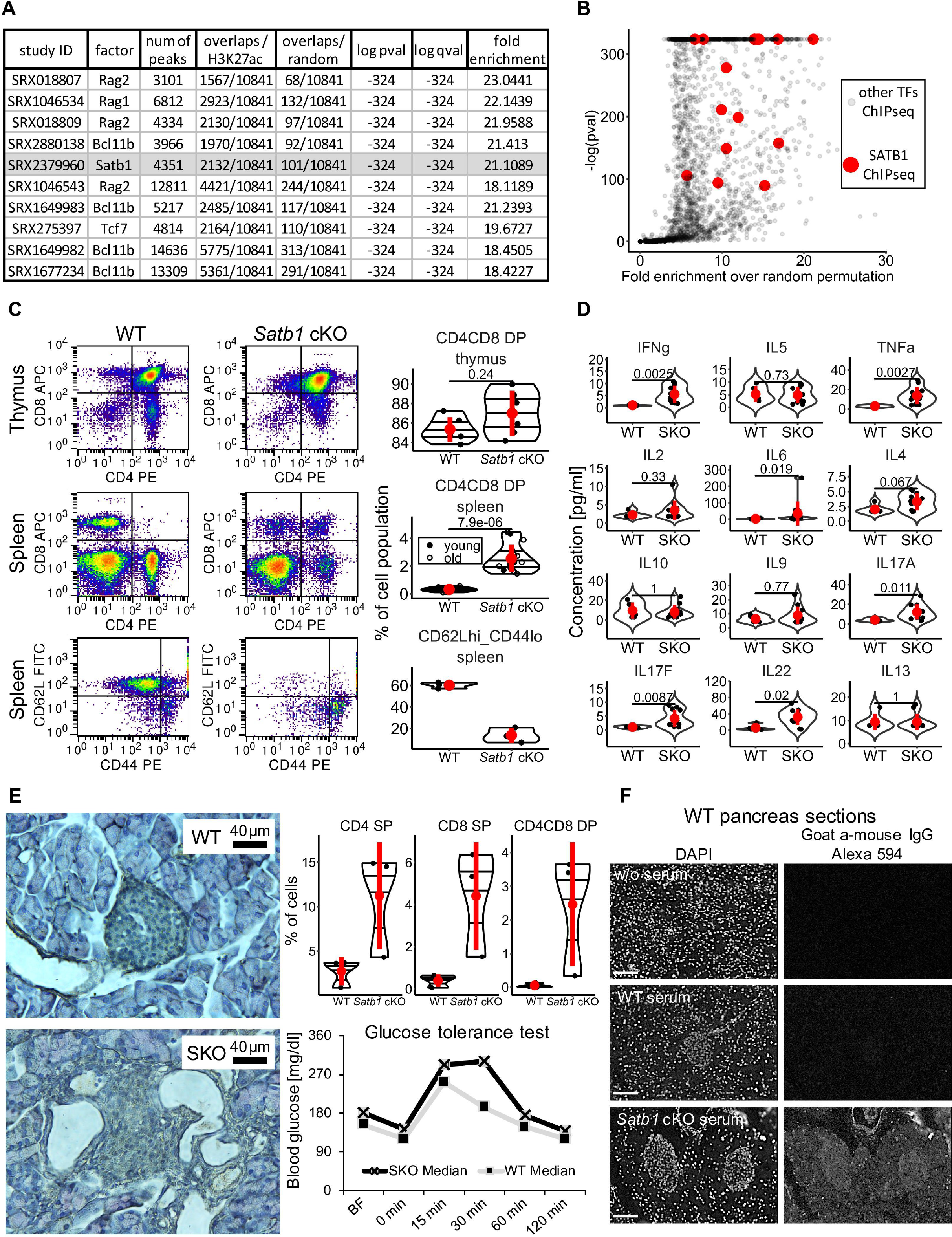
**Autoimmune-like phenotype of the Satb1 cKO mouse** (A) Anchors of regulatory H3K27ac HiChIP loops in murine WT thymocytes are enriched for the ChIP-seq peaks of the factors depicted in the table. The analysis was based on the ChIP-Atlas (Oki et al., 2018), thus the datasets and underlying study ID match SRA databases. (B) Distribution of fold-enrichment of all available ChIP-seq datasets based on ChIP-Atlas (Oki et al., 2018) at WT H3K27ac loop anchors over random permutation and the corresponding p values. All available SATB1 ChIP-seq datasets (highlighted in red) evinced enrichment at the anchors of regulatory chromatin loops. (C) Flow cytometry in cell populations of the thymus and the spleen for the expression of CD4 and CD8 cell surface markers as well as CD62L (naiveness marker) and CD44 (lymphocyte activation marker). Used animals were divided into young (45±11 days; 6 WT, 6 Satb1 cKO) and old (179±35 days; 7 WT, 9 *Satb1* cKO) age categories. Only young animals were used for analysis of thymus due to its deterioration in old animals. Three young animals for each genotype were used for CD62 / CD44 analysis. (D) Differences in the cytokine milieu in the blood serum of WT and *Satb1* cKO animals measured with bead-based immunoassay, point to an elevated Th17 response and increased inflammatory cytokines. (E) *Satb1* cKO animals display an autoimmune-like phenotype. T cells infiltrated peripheral organs, including pancreas, which resulted into damaged islets of Langerhans and consequently impaired glucose metabolism. BF indicates steady-state glucose levels, before 6 hour fasting period. (F) WT pancreas sections were incubated with serum from either WT or *Satb1* cKO animals to detect the presence of autoantibodies. Scale bar in all images is 100 μm. In (C, D and E) if present, the horizontal lines inside violins represent the 25^th^, 50^th^ and 75^th^ percentiles. Red circle represents mean ± s.d. *P* values by Wilcoxon rank sum test. In (D and E), SKO represents *Satb1*^fl/fl^*Cd4-*Cre^+^ animals.

### The ablation of SATB1 from murine T cells leads to autoimmunity

In order to link the molecular mechanisms governing T cell chromatin organization to physiology, we generated a *Satb1*^fl/fl^*Cd4-*Cre^+^ (*Satb1* cKO) conditional knockout mouse and characterized its phenotype. The knockout animals displayed problems with their skin and fur, inflammation in various tissues and affected lymphoid organs (Figure S1A). The thymi of *Satb1* cKO animals were smaller in size, unlike their enlarged peripheral lymphoid organs. Thymic deregulation was also reflected in the impaired developmental processes in the thymus as demonstrated by the deregulation of T cell populations (Figure 1C and S1B). The increased number of CD4^+^CD8^+^ (double positive - DP) cells and the decreased numbers of CD4^+^ and CD8^+^ single positive (SP) cells in the thymus of *Satb1* cKO mice indicated a developmental blockade at the DP stage, pointing to altered positive selection as previously suggested (Alvarez et al., 2000; Kondo et al., 2016). Moreover, there was a diminished pool of naïve CD62L^hi^CD44^lo^ peripheral T cells and an increased fraction of CD44^hi^ T cells displaying an activated (and/or memory) T cell phenotype (Figure 1C). Deregulation of the thymic developmental programs was also supported by the altered cytokine milieu in the blood serum, with prevailing IL-17 cell responses and increased levels of pro-inflammatory cytokines such as IFNγ and TNFα detected in *Satb1* cKO sera (Figure 1D). The increased levels of DP T cells in the spleen (Figure 1C), together with the absence of naïve CD4^+^ T cells were suggestive of an autoimmune-like phenotype (Sakaguchi et al., 2008). Indeed, we observed infiltration of T cells in the pancreas of the *Satb1* cKO animals, causing damage to the islets of Langerhans and leading to impaired glucose metabolism (Figure 1E). The deregulation of cellular immunity was accompanied by the presence of autoantibodies, which we demonstrated by incubating sections of WT pancreas with *Satb1* cKO sera (Figure 1F) and *Satb1* cKO sections of pancreas and lungs with *Satb1* cKO sera (Figure S1C). Based on these findings, we concluded that SATB1 absence leads to impaired thymocyte development and the concomitant deregulation of T cell populations in the secondary lymphoid organs, affecting T cell homeostasis and sustaining an autoimmune-like phenotype.

### Roles of SATB1 and CTCF in T cell chromatin organization

SATB1 has been previously attributed with genome organizing functions (Cai et al., 2003, 2006), therefore we aimed to investigate the potential deregulation of thymocyte genome organization that is anticipated upon SATB1 depletion and link it to the deregulation of immune physiology we observed in the *Satb1* cKO mice. For this purpose, we performed Hi-C experiments (Lieberman-Aiden et al., 2009) in both WT and *Satb1* cKO thymocytes (Table S1). We did not identify any major changes in high-order chromatin organization (Figure 2A). Differential analysis of topologically associating domains (TADs) between WT and *Satb1* cKO cells also supported this claim with an average of 77% unchanged TADs between WT and *Satb1* cKO, resembling the level of differences being usually detected between the different biological replicates of the same experiment (Figure S1D; Dixon et al., 2012; Rao et al., 2014; Sauerwald et al., 2020). Even though broad scale differences were not observed in the Hi-C maps, one may interrogate more localized conformational changes with HiChIP data, especially given our underlying hypothesis of transcription factor-guided genome organization. Therefore, we next compared the SATB1-mediated and CTCF-mediated chromatin loops by performing HiChIP experiments targeting the respective factors in WT cells. Our HiChIP datasets at 5 kbp resolution (FDR ≤ 0.01) yielded 1,374 and 3,029 loops for SATB1 and CTCF, respectively (Table S1, S2). It is important to note that in the SATB1 HiChIP experiments we used custom-made antibodies specifically targeting the long SATB1 isoform that we recently characterized (Zelenka et al., Submitted; Table S3). We compared differentially interacting areas of the HiChIP matrices at 100 kbp and 500 kbp resolution (Figure 2B). At 100 kbp resolution, 46 interaction pairs were stronger in the SATB1 contact matrix compared to 553 in the CTCF matrix (FDR ≤ 0.05). The analysis at 500 kbp resolution indicated a similar disproportion (7 vs 42), collectively suggesting that CTCF contributes to the high-order chromatin organization in developing T cells to a much higher extent than SATB1. Next, we performed aggregate peak analysis (APA; Rao et al., 2014) applying the SATB1/CTCF-mediated HiChIP loops on Hi-C datasets derived from WT and *Satb1* cKO thymocytes. As expected, this analysis unraveled diminished interactions for the SATB1-mediated loops in the *Satb1* cKO cells compared to WT, but no change was evident for the CTCF-mediated loops (Figure S1E). Together with the unchanged RNA levels of *Ctcf* in the *Satb1* cKO, this suggested that CTCF was capable of maintaining the high-order chromatin structure in the *Satb1* cKO cells. Moreover, genes residing in both CTCF- and SATB1-mediated loops were transcriptionally insulated from their gene neighbors (Figure S1F) and this characteristic was not altered in the *Satb1* cKO. The latter was not surprising since out of 1,374 SATB1-mediated loops, the vast majority (84%) overlapped with at least one CTCF-mediated loop (Figure 2C). An overlap score, calculated by dividing the length of the overlap by the total size of a loop, indicated that most of SATB1-mediated loops were engulfed within CTCF loops (Figure 2D). As CTCF is a well-characterized protein with insulator function (Phillips and Corces, 2009), it is likely that the transcriptional insulation effect of SATB1 was derived from the CTCF function. Nevertheless; the binding pattern of these factors was quite different. Similar to previously published results (Ghosh et al., 2019), the SATB1 binding sites we have identified, evinced a nucleosome preference, unlike CTCF (Figure 2E). Gene ontology analysis of the genes intersecting with loop anchors uncovered the high propensity of SATB1 to participate in the loopscape structure of immune-related genes, while CTCF-mediated chromatin loops exhibited omnipresent looping patterns resulting in the enrichment of general metabolic and cellular processes (Figure 2F). Taking these results under consideration we conclude that the high-order chromatin organization of murine thymocytes is primarily maintained via CTCF long-range chromatin interactions with minor input from the SATB1-mediated loops.

**Figure 2.**
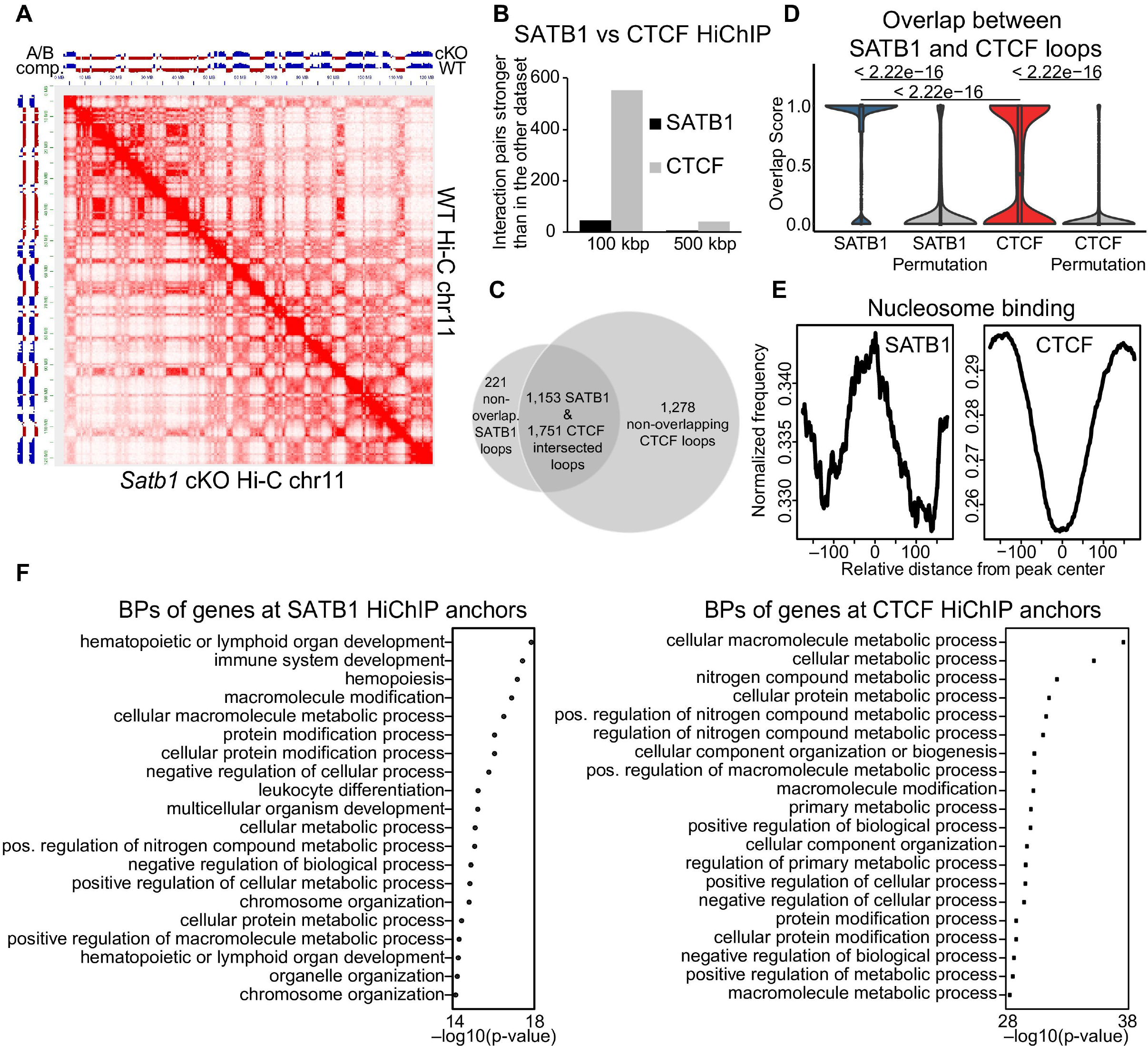
**T cell chromatin organization mediated by SATB1 and CTCF** (A) Comparison of WT and *Satb1* cKO Hi-C heatmaps of chromosome 11 indicates no major changes at high order chromatin level in murine thymocytes. (B) diffHic analysis (Lun and Smyth, 2015) of differentially interacting chromatin areas indicates that CTCF contributes more strongly to the higher order chromatin organization of the murine T cell genome than SATB1. (C) SATB1-mediated loops highly intersect with CTCF-mediated loops detected by HiChIP. For the intersection, the outer coordinates of left and right loop anchors were used. (D) Overlap score between SATB1 and CTCF loops calculated as (number of overlapping bp) / (bp size of a loop). For example, for the SATB1-labeled violin, a score of 1.0 indicates either 100% overlap or engulfment of a SATB1-mediated loop in a loop mediated by CTCF. A score of 0.0 indicates no overlap. The plot indicates that the majority of the SATB1 loops were engulfed in CTCF loops. The same approach was repeated for randomly shuffled loops. *P* values by Wilcoxon rank sum test. (E) SATB1 preferentially binds nucleosomes unlike CTCF. (F) SATB1 loop anchors overlap with genes enriched for immune system-related categories. CTCF-mediated loops display more widespread coverage of intersecting genes thus the most enriched gene ontology pathways belong mostly to general cellular processes.

### The regulatory role of SATB1-mediated chromatin loops in murine T cells

To unravel the regulatory potential of SATB1-mediated chromatin loops we investigated the impact of SATB1 depletion in *Satb1* cKO thymocytes. These cells generally appeared to have more compact chromatin as demonstrated by several measures. Immunofluorescence experiments displayed more intense HP1α staining (marker of heterochromatin) in *Satb1* cKO thymocytes (Figure 3A). Despite the gross similarities at the higher order chromatin structure between WT and *Satb1* cKO cells, as deduced by Hi-C, we have detected 1.11% of chromosome compartments that turned from compartment A (predominantly consisting of euchromatin) to compartment B (heterochromatin regions; Lieberman-Aiden et al., 2009) in the *Satb1* cKO cells, compared to 0.59% of B to A compartment switch (Figure S1G). Genes affected by the aforementioned A-to-B compartment switch did not display any gene ontology pathway enrichment which would otherwise be indicative of a link between high-order chromatin structure and the deregulated immune system in *Satb1* cKO animals. This observation further reinforced our hypothesis that SATB1 acts at a finer-scale level of genome organization. In addition, *Satb1* cKO cells evinced a higher fraction of less accessible regions (6,389 compared to 5,114 more accessible regions; p ≤ 0.01) based on ATAC-seq analysis performed for WT and *Satb1* cKO thymocytes (Figure S1H; Table S4). To determine whether these chromatin accessibility changes were also reflected at the transcriptional level, we performed stranded total RNA sequencing. Our analysis revealed that 922 genes were significantly underexpressed and 719 genes were significantly overexpressed in the *Satb1* cKO compared to WT thymocytes (FDR<0.05; Table S5). Such a strong deregulation of the transcriptional landscape in *Satb1* cKO cells in contrast to the modest transcriptional changes observed upon depletion of conventional genome organizers (Nora et al., 2017; Rao et al., 2017; Schwarzer et al., 2017) emphasizes the regulatory importance of SATB1-dependent chromatin organization.

**Figure 3.**
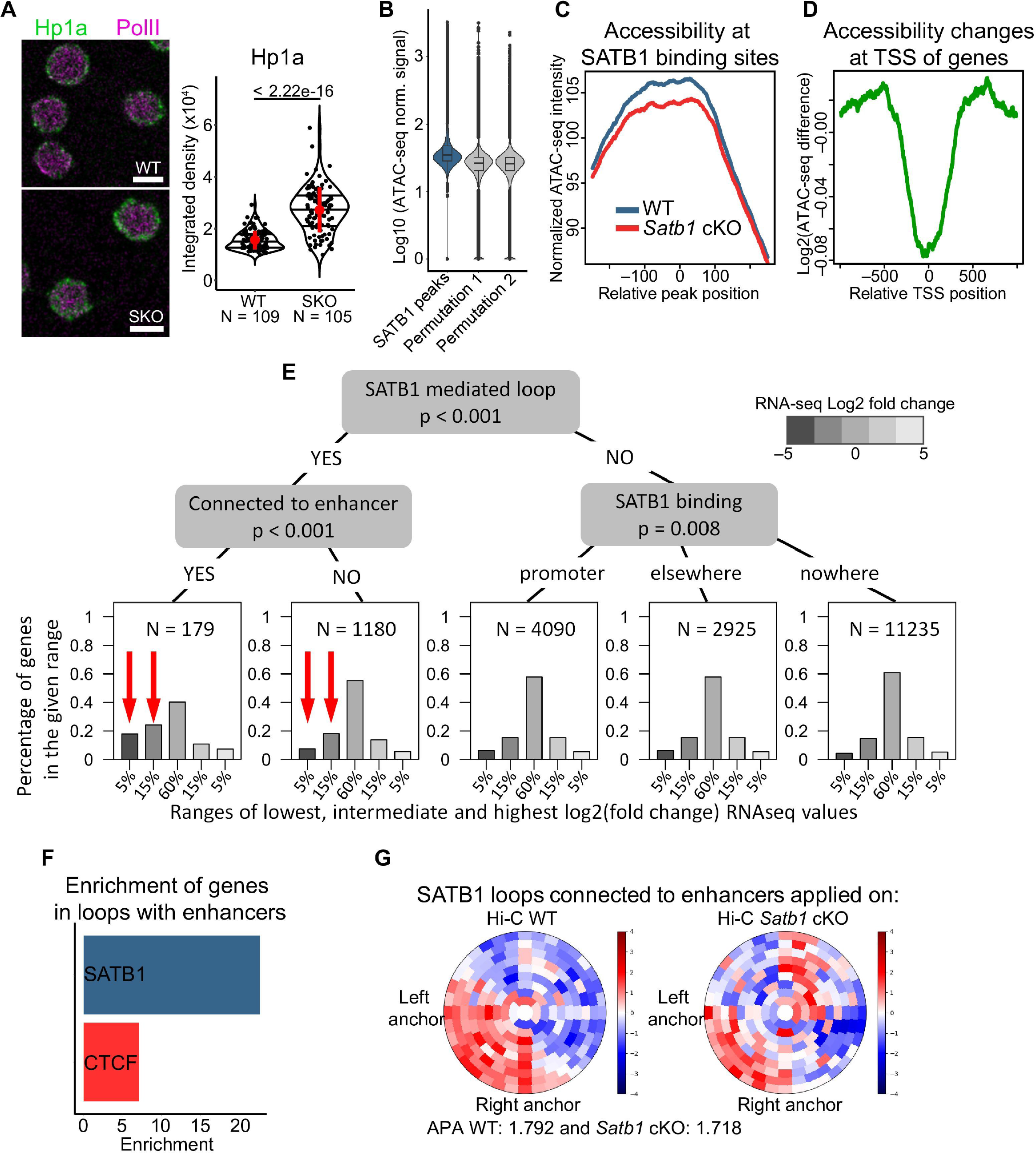
**SATB1-mediated chromatin loops positively regulate gene expression** (A) Immunofluorescence analysis with confocal microscopy of WT and *Satb1* cKO thymocytes stained with antibodies against HP1α and RNA Pol II. The nuclei of *Satb1* cKO thymocytes had stronger HP1α signal, suggesting a more repressed nuclear environment. The values in the graph represent an integrated signal density from summed z-stacks. SKO represents *Satb1*^fl/fl^*Cd4-*Cre^+^ animals. The horizontal lines inside violins represent the 25^th^, 50^th^ and 75^th^ percentiles. The red circle represents the mean ± s.d. *P* values by Wilcoxon rank sum test. Scale bar 5 μm. (B) ATAC-seq signal indicates higher chromatin accessibility at WT SATB1 binding sites than expected by chance (i.e, randomly shuffled SATB1 binding sites). 100 randomizations were used to for statistical evaluation (bootstrap p-value = 0). Two representative random distributions are depicted in the figure. (C) Chromatin accessibility at SATB1 binding sites is decreased in *Satb1* cKO. (D) Log2 fold change of chromatin accessibility indicates the highest accessibility drop in *Satb1* cKO being at the TSS of genes. (E) Inference tree systematically probing all options of SATB1 binding and looping as well as their impact on gene expression. The x-axis indicates the ranges of log2 fold change RNA-seq values. For example, 5% of the most underexpressed genes in *Satb1* cKO represent ∼20% of genes (y-axis) found in anchors of SATB1-mediated loops connected to an enhancer (first red arrow). The red arrows highlight the disruption of the normal distribution of differentially expressed genes in the respective ranges that is present in SATB1-mediated loops. Differentially expressed genes that do not overlap with anchors of SATB1 loops display a normal distribution – reflecting the respective log2 fold change ranges (the three rightmost graphs). (F) SATB1-mediated loops connecting genes to enhancers are about three-fold enriched compared to CTCF loops. (G) SATB1-mediated loops connected to enhancers evince enriched interaction signal between left and right anchor in WT Hi-C data, which is deregulated in *Satb1* cKO. Aggregate peak analysis (APA; Rao et al., 2014) was calculated and visualized by SIPMeta (Rowley et al., 2020).

SATB1 is known for its functional ambiguity of acting either as a transcriptional activator or a repressor, depending on the cellular context (Kumar et al., 2006). Although the original studies were mainly focused on its repressive roles (Kohwi Shigematsu et al., 1997; Liu et al., 1997; Seo et al., 2005), our aforementioned observations supported the increased chromatin compactness and subsequent repressed environment of the SATB1-depleted cells. This rather indicated its positive impact on transcriptional gene regulation. The vast majority of SATB1 binding sites in WT thymocytes evinced increased chromatin accessibility compared to randomly shuffled binding sites (i.e. what would be expected by chance, 100 randomizations used, bootstrap p-value = 0; Figure 3B), with a visible drop in chromatin accessibility in the *Satb1* cKO (Figure 3C). This drop in chromatin accessibility in *Satb1* cKO cells was especially evident at the transcription start site of genes (TSS), suggesting a direct role of SATB1 in gene transcription regulation (Figure 3D and S1I). Moreover, the expression changes of significantly deregulated genes were positively correlated with the changes of chromatin accessibility at promoters determined by ATAC-seq (Spearman’s ρ = 0.438, *P* < 2.22e–16; Figure S1J). Only about 5% of SATB1 binding sites had low chromatin accessibility in WT (lower than the average accessibility score of ten randomizations depicted in Figure 3B). These regions had increased chromatin accessibility in the *Satb1* cKO (Figure S1K), which would suggest a repressive function of SATB1. However, these regions were not enriched for immune-related genes (not shown) and thus probably not contributing to the observed phenotype.

We have next created a linear regression model, as an unbiased way to identify how gene expression was affected in murine thymocytes upon SATB1 depletion. We utilized SATB1 binding, SATB1- and CTCF-mediated chromatin loops, changes in H3K27ac-dependent chromatin loops and changes in chromatin accessibility at different positions of a gene, as predictors of RNA level changes between WT and *Satb1* cKO cells. We found that the majority of predictors exhibited an expected behavior, such as increased chromatin accessibility at gene promoters was associated with increased gene expression (Figure S2). The regression model highlighted SATB1 binding and SATB1-mediated chromatin loops as good predictors associated with decreased RNA levels of influenced genes in the *Satb1* cKO. In this analysis, we applied the model for all known genes, which resulted in a quite low R-squared value (0.113). For this reason, we verified the activatory role of SATB1 loops with an additional approach. We constructed an inference tree, where we systematically probed the distribution of gene expression for the genes that were or were not found in SATB1-mediated loops (Figure 3E). Genes located in SATB1-mediated loops displayed lower RNA levels in the *Satb1* cKO, an effect that was further intensified when the gene was connected to an enhancer (Figure 3E), suggesting a positive role for SATB1 in gene transcription via promoter-enhancer mediated chromatin loops.

Thymic enhancers were previously shown to be occupied by conventional genome organizers such as CTCF and cohesin, suggesting their involvement in gene regulatory loops of thymocytes (Ing-Simmons et al., 2015; Seitan et al., 2013). We utilized a list of predicted thymic enhancers (Shen et al., 2012) and we found more than 2-fold enrichment of SATB1-over CTCF-mediated loops overlapping with such enhancers (Figure S1L) and more than 3-fold enrichment of genes connected to enhancers by SATB1-mediated chromatin loops over CTCF-mediated loops (Figure 3F). SATB1-mediated loops connected to enhancers, also displayed a disturbed chromatin interaction pattern in *Satb1* cKO Hi-C data compared to WT (Figure 3G). Collectively, these findings suggested that CTCF participates in mechanisms responsible for supporting a basal high-order T cell chromatin structure, whereupon SATB1 likely exerts its action in a more refined organization layer consisting of promoter-enhancer chromatin loops.

### Deregulated promoter-enhancer loops in *Satb1* cKO T cells

To further investigate the latter hypothesis, we compared the promoter-enhancer chromatin loops present in WT and *Satb1* cKO thymocytes, utilizing the H3K27ac HiChIP loops. H3K27ac HiChIP in *Satb1* cKO thymocytes yielded 19,498 loops (compared to 16,458 loops detected in WT; Table S1, S2). Differential analysis of the 3D interactions (independent on the 1D H3K27ac ChIP-seq signal) identified 11,540 and 12,111 H3K27ac loops displaying decreased or increased contact enrichment in the *Satb1* cKO compared to WT cells, respectively (further referred as “underinteracting” and “overinteracting” H3K27ac loops, respectively; Table S6). The RNA levels of the genes associated with differential H3K27ac chromatin loops in WT versus *Satb1* cKO displayed a positive correlation (Spearman’s ρ = 0.26) with the difference between over- and under-interacting H3K27ac loops (Figure 4A). The SATB1-mediated underinteracting H3K27ac loops displayed the highest drop in the RNA levels of the overlapping genes compared to those in non-SATB1 underinteracting H3K27ac loops (Figure 4B). In contrast, the genes localized in anchors of overinteracting H3K27ac loops did not show any major changes in expression (Figure 4C). Moreover, the expression of genes located at anchors of SATB1-mediated loops was decreased more dramatically than genes located at CTCF loops (Figure 4B). This finding is a good indication of causality, supporting direct involvement of SATB1 in the regulatory chromatin loops.

**Figure 4.**
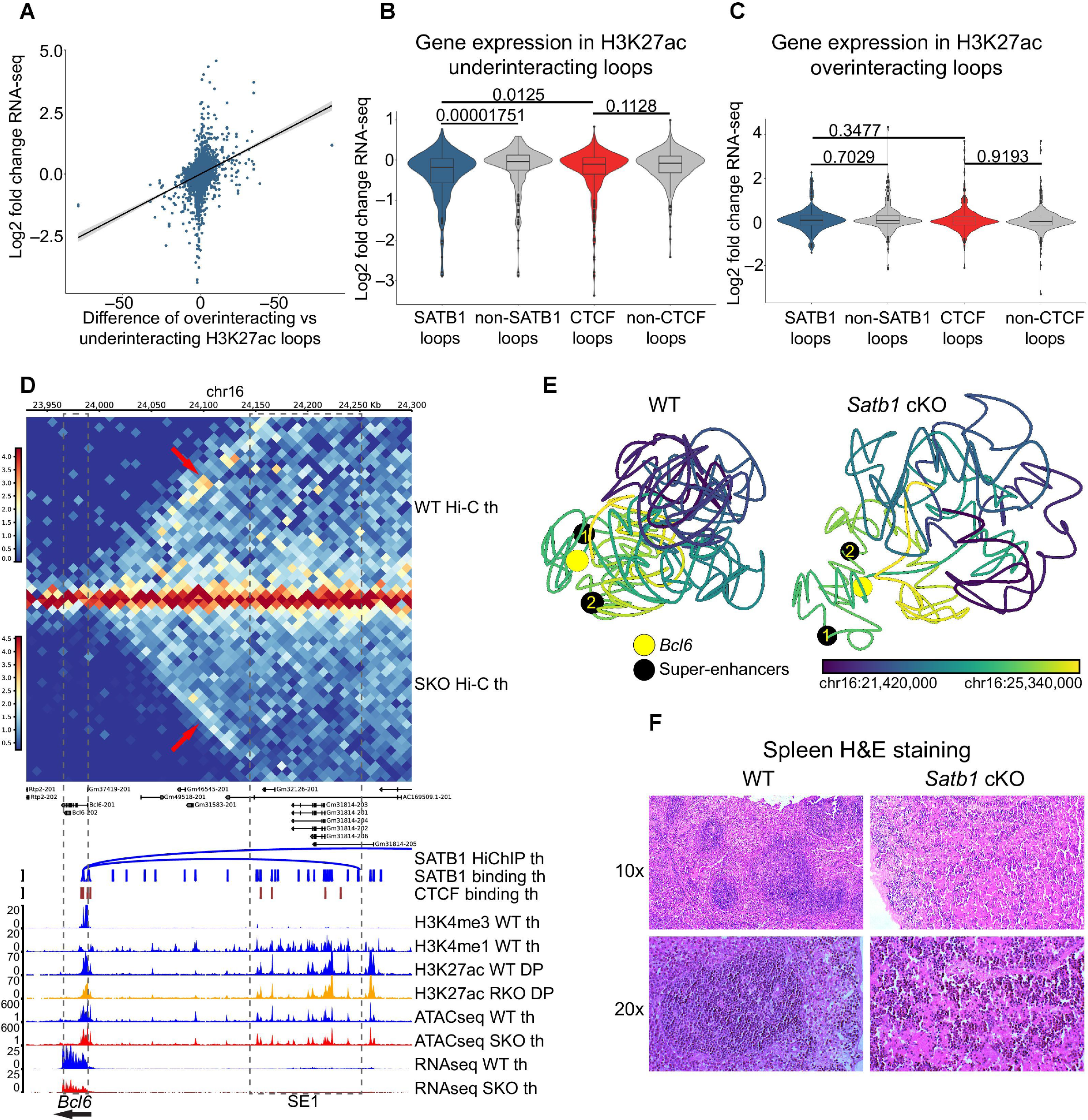
**SATB1 mediates promoter-enhancer communication of critical immune-related genes** (A) Positive correlation (Spearman’s ρ = 0.26) between gene expression changes and the difference between H3K27ac overinteracting and underinteracting loops in WT thymocytes compared to *Satb1* cKO. Negative values on the x-axis represent prevailing H3K27ac loops that were lost or diminished in *Satb1* cKO, whereas positive numbers refer to gained H3K27ac loops in the knockout. (B) Log2 fold change expression values of genes that were present in anchors of H3K27ac HiChIP loops underinteracting in *Satb1* cKO and which also did or did not intersect with SATB1/CTCF-mediated loops. An equal number of underinteracting loops that did not overlap with SATB1/CTCF-mediated loops were randomly generated. *P* values by Wilcoxon rank sum test. (C) The same as in (B) visualized for H3K27ac HiChIP loops overinteracting in *Satb1* cKO. (D) The *Bcl6* gene is connected by two SATB1 loops to its super-enhancer regions. Figure depicts the shorter SATB1 loop connecting *Bcl6* (same loop anchor for both loops; chr16:23985000-23990000) with the more proximal super-enhancer region 1 (SE1; chr16:24245000-24250000) and a part of the larger SATB1 loop (chr16:24505000-24510000). In *Satb1* cKO, these interactions were lost as seen in the Hi-C data analysis (heatmap in the top). Legend: th – thymocytes, DP – CD4^+^CD8^+^ T cells, RKO – *Rad21*^fl/fl^*Cd4-*Cre^+^ and SKO – *Satb1*^fl/fl^*Cd4-*Cre^+^. (E) Computational 3D modeling utilizing the WT Hi-C and *Satb1* cKO Hi-C data, to visualize the proximity between the *Bcl6* gene body and its two super-enhancer regions / SATB1 loop anchors. The black beads represent the edge of super-enhancer regions demarcated by the SATB1 loop anchors [SE1 and the short loop are also depicted in (D)]. Color gradient represents linear genomic position along the locus. (F) Hematoxylin-Eosin staining of WT and *Satb1* cKO spleen sections revealed disturbed germinal centers in the knockout.

Next, we sought to investigate the genes intersecting with anchors of the differential H3K27ac loops. Underinteracting H3K27ac loops, with the highest score, included genes encoding for master regulators and T cell signature genes such as *Bcl6*, *Ets2*, *Tcf7*, *Cd8b1*, *Ikzf1*, *Bach2*, *Cd6*, *Rag2*, *Il4ra*, *Rag1*, *Lef1* and others (in descending order; Table S6), which all evinced decreased RNA levels in the *Satb1* cKO. On the other hand, the overinteracting H3K27ac loops with the highest score also contained factors essential for proper T cell development and differentiation such as *Tox*, *Gata3*, *Ifngr1, Maf* and/or *Jun* (Table S6), which all correspondingly displayed increased gene expression in the *Satb1* cKO thymocytes. Certain genes present in the overinteracting H3K27ac loops were bound by SATB1 and a fraction of them were also found in SATB1-mediated loops. Thus, we cannot exclude the possibility that SATB1 mediates a repressive role for these targets. Though, since our unbiased approaches have primarily suggested an activatory role for the SATB1-mediated loops, in this work we focused on this.

### SATB1 positively regulates *Bcl6* and other master regulator genes

The most highly affected candidate gene, in terms of H3K27ac underinteracting loops in the *Satb1* cKO compared to WT, was *Bcl6*. The expression of *Bcl6* gene in B cells is regulated by a set of super-enhancers; one spanning the promoter and 5’ UTR region, and additional three distal upstream enhancer stretches at 150-250 kbp, ∼350 kbp and at ∼500 kbp (Chapuy et al., 2013; Qian et al., 2014; Ramachandrareddy et al., 2010; Ryan et al., 2015). Apart from H3K27ac differential loops, we identified increased SATB1-mediated interactions in the locus with two significant SATB1-mediated loops connecting *Bcl6* and the super-enhancer regions at ∼250 kbp and ∼500 kbp upstream of the gene (Figure 4D; here referred to as SE1 and SE2, respectively). These enriched chromatin interactions observed in WT were absent in the *Satb1* cKO thymocytes as deduced by Hi-C experiments (red arrows). As a result of this deregulation, the *Bcl6* gene displayed significantly lower RNA levels in the *Satb1* cKO thymocytes (log2FC = –1.290, FDR = 5.6E–10). Next, we utilized this gene locus as an example for 3D modeling experiments. Initially, we utilized our CTCF and SATB1 HiChIP data and performed computational modeling which supported the idea that SATB1-mediated chromatin landscape represented a regulatory layer, built on a generic scaffold mediated by other factors, at least partly by CTCF (Figure S3A). 3D modeling based on Hi-C data allowed us to better visualize the differences in the proximity between *Bcl6* and its super-enhancers in WT and *Satb1* cKO cells (Figure 4E). To further support the functional significance of these interactions, we overlaid these models with ChIP-seq data for the histone modifications H3K27ac, H3K4me1 and H3K4me3 from WT cells (Figure S3B). These models showed that active enhancers decorated by H3K27ac and H3K4me1 were located in spatial proximity to *Bcl6* gene in WT and not in *Satb1* cKO cells. It is worth noting that 1D H3K27ac ChIP-seq peaks derived from HiChIP experiments available for WT and *Satb1* cKO did not reveal any major differences between the genotypes, which further reinforces the importance of SATB1-mediated 3D chromatin organization regulating *Bcl6* expression. 3D modeling, based on thymocyte Hi-C datasets, allowed us to untangle the potential presence of discrete cell subpopulations. In WT animals, we identified two subpopulations of cells differing in the proximity between *Bcl6* and its super-enhancers (Figure S3C); yet no significant subpopulation formation was detectable in the *Satb1* cKO. BCL6 is the master regulator of Tfh cell lineage specification during the differentiation of naive CD4 cells into Tfh cells (Johnston et al., 2009; Nurieva et al., 2009; Yu et al., 2009). However, it is also expressed in developing thymocytes (Hyjek et al., 2001; Sun et al., 2000) where it was shown to form a complex with E3 ubiquitin ligase CUL3 and exert a negative feedback loop on the Tfh program by repressing *Batf* and *Bcl6* (Mathew et al., 2014). Thus, we speculated that the different subpopulations identified in our 3D modeling may be linked to the different developmental T cell fates, where Tfh precursor cells would differ from other cell type precursors by the distance between *Bcl6* and its super-enhancers. In support to the connection between SATB1-mediated regulation and Tfh lineage specification, we demonstrated that *Satb1* cKO animals had disturbed germinal centers (Figure 4F), which translated into the production of autoantibodies (Figure 1F and S1C).

Apart from *Bcl6*, more genes (depicted in Table S6) were also regulated by the SATB1-mediated chromatin organization which are quite important for the function of the immune system. Genomic tracks and SATB1-mediated HiChIP loops for such selected genes (*Tcf7*, *Lef1*, *Cd8*, *Ikzf1*, *Satb1*) are presented in Figure S4.

### T cell receptor locus and cell adhesion in *Satb1* cKO

We should note that in the differential analysis of H3K27ac loops, short genes could be underestimated, hence we further considered gene length in our analysis. Indeed, upon taking this into account, the most affected genes in both categories of overinteracting and underinteracting loops were enriched for gene segments of the T cell receptor (TCR) locus (Table S6).

TCR is the most important cell surface receptor expressed in thymocytes which defines multiple developmental decisions. The generation of a functional TCR involves recombination of the variable (V), diversity (D) and joining (J) gene segments via a process called V(D)J recombination. This process is based upon the action of protein complexes that include RAG1 and RAG2 recombinases (Fugmann et al., 2000). Recruitment of RAG proteins is highly correlated with active promoters labeled with H3K4me3 (Ji et al., 2010; Teng et al., 2015) and also with enhancers and regions decorated by the H3K27ac mark (Maman et al., 2016; Teng et al., 2015). Moreover, recombination is regulated by the 3D organization of the locus driven by the architectural proteins CTCF and cohesin, mainly via the arrangement of specific TCR enhancers (Chen et al., 2015; Seitan et al., 2011; Shih et al., 2012).

Here, we revealed a number of SATB1-mediated loops connecting the TCRα enhancer with inner regions of the locus (Figure 5A). Moreover; a number of highly significant SATB1 loops split the region of joining gene segments into two parts – one part containing gene segments that were overexpressed and the other half containing gene segments that were underexpressed in the *Satb1* cKO. This resulted in the defective usage of the TCRα joining segments (Figure 5A), coupled with the overall *Tcra* rearrangement in *Satb1* cKO animals as previously reported (Feng et al., 2021; Hao et al., 2015). A previous study ascribed this deregulation to the lost SATB1-mediated regulatory loops, positively controlling the expression of both *Rag1*/*Rag2* genes resulting in lower levels of RAG proteins (Hao et al., 2015). We validated the presence of these regulatory loops (Figure 5B) as well as the resulting 2-fold and 2.8-fold decrease in thymic RNA levels of the *Rag1* and *Rag2* genes (Figure 5C), respectively (which was less profound in sorted DP cells: 1.34-fold and 1.61-fold decrease for *Rag1* and *Rag2* genes, respectively; data not shown). However, given the long turnover of RAG proteins, their thymic protein levels were not significantly affected (Figure 5D). Moreover, the representation of *Traj* fragments was correlated with the presence of overinteracting and underinteracting H3K27ac loops (Figure 5E), indicating the importance of 3D chromatin organization of the TCR locus in its rearrangements.

**Figure 5.**
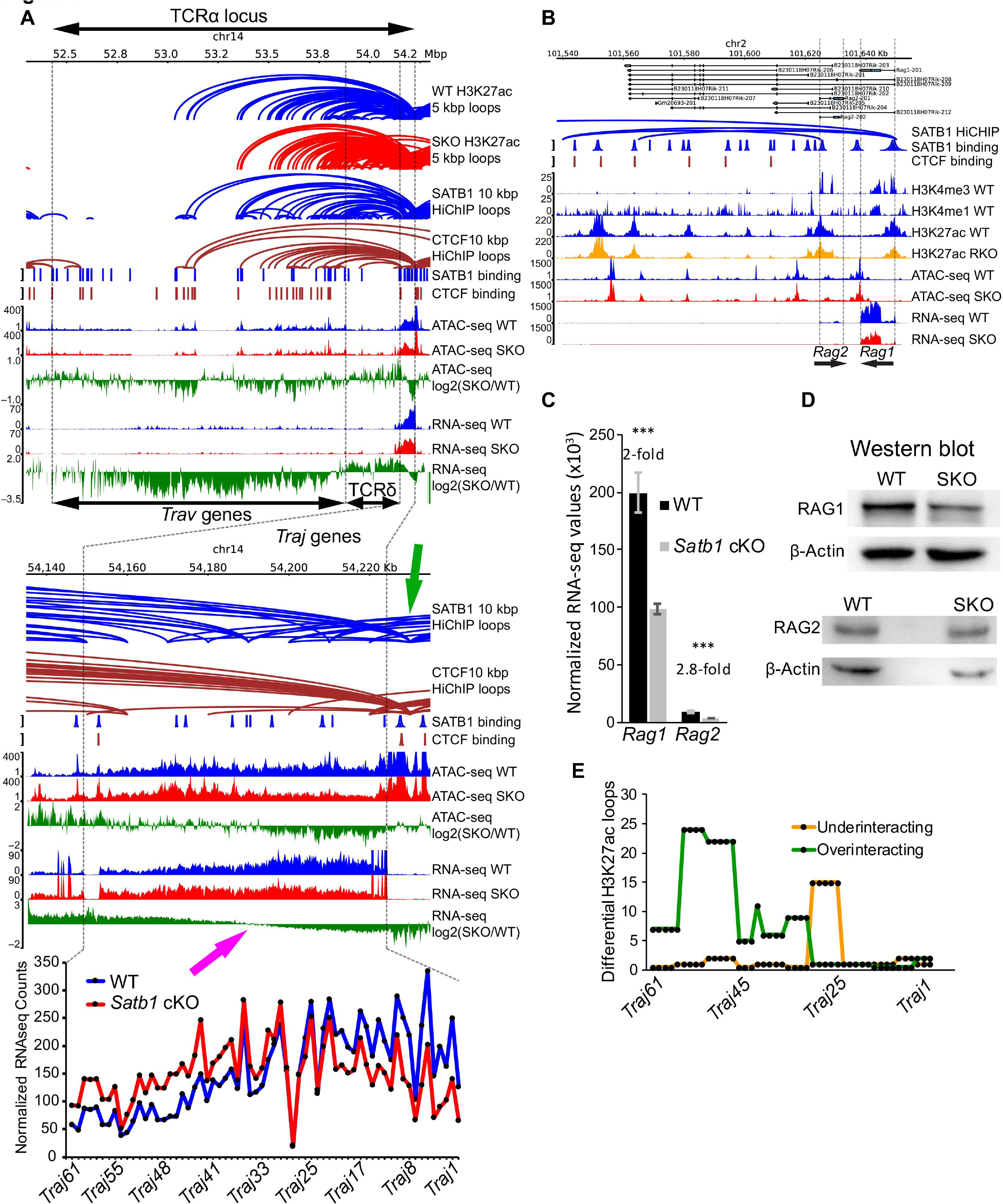
**TCRα gene locus chromatin organization depends on SATB1** (A) Genomic tracks as well as SATB1 and CTCF HiChIP loops at the T cell receptor alpha locus (TCRα). The bottom green genomic tracks of log2 fold change RNA-seq values summarize the overall deregulation of the TCRα locus, with variable regions (*Trav* genes) being mostly underexpressed, TCRδ locus overexpressed and TCRα joining regions (*Traj* genes) displaying geometrically symmetric deregulation splitting the region into over- and under-expressed in *Satb1* cKO cells (magenta arrow). This deregulation was markedly correlated with the presence of SATB1-mediated loops. Note especially the region of joining genes which manifests a deregulation similar to previous reports from *Satb1* depleted animals (Feng et al., 2021; Hao et al., 2015). Both SATB1 and CTCF loops displayed a tendency to connect the TCR enhancer (green arrow) to the regions inside the locus. The presented loops were called with low stringency parameters and with a different set of binding sites compared to the rest of our study due to technical details explained in the methods section. (B) SATB1 mediates promoter-enhancer regulatory loops controlling the expression of both *Rag1* and *Rag2* genes. (C) RNA levels of *Rag1* and *Rag2* genes in WT and *Satb1* cKO thymocytes. (D) Protein levels of RAG1 (1:2500, Abcam ab172637) and RAG2 (1:100, Santa Cruz Biotechnology sc-517209) were just marginally affected in the *Satb1* cKO thymocytes (beta Actin as a loading control; 1:500, ORIGENE TA811000). (E) Differential H3K27ac HiChIP loops reflected the deregulation of *Traj* regions. In (A, B and D), H3K27ac datasets originate from CD4^+^CD8^+^ T cells, all the other datasets from thymocytes. RKO denotes *Rad21*^fl/fl^*Cd4-*Cre^+^ and SKO – *Satb1*^fl/fl^*Cd4-*Cre^+^.

Apart from the deregulation of H3K27ac chromatin loops in *Satb1* cKO compared to WT thymocytes, as deduced by HiChIP experiments we should also note that the “cell adhesion pathway” was overrepresented in gene ontology analysis for both underexpressed genes by RNA-seq and genes associated with less accessible regions as deduced by ATAC-seq (Figure 6A; gene ontology pathways for other genomic datasets are presented in Figure S5). Indeed, genes encoding key molecules for intrathymic crosstalk (Lopes et al., 2015) were underexpressed in the *Satb1* cKO (Figure 6B). In addition to the deregulated loopscape structure of the TCRα gene locus, we identified similar defects in the SATB1-mediated regulatory looping for adhesion molecule gene loci such as *Cd28* (Figure 6C), *Lta*, *Ltb* (Figure S6A) and *Ccr7* (Figure S6B). Total thymocyte RNA sequencing revealed the downregulation in expression levels of receptors specific for the medullary thymic epithelial cells (Figure 6B). In support of this finding, we also identified the disrupted thymic structure and impaired cell-to-cell communication in the *Satb1* cKO thymus. This deregulation was notable from several histological and electron microscopy experiments indicating disrupted cellular contacts in *Satb1* cKO (Figure 6D). As a result, the thymi of *Satb1* cKO animals contained a lower number of cells (Figure 6E), partly due to the increased apoptosis rate (Figure 6F) resulting from the impaired developmental pathways and partly due to the increased exit rate of these improperly developed T cells from the thymus (Figure 6G).

**Figure 6.**
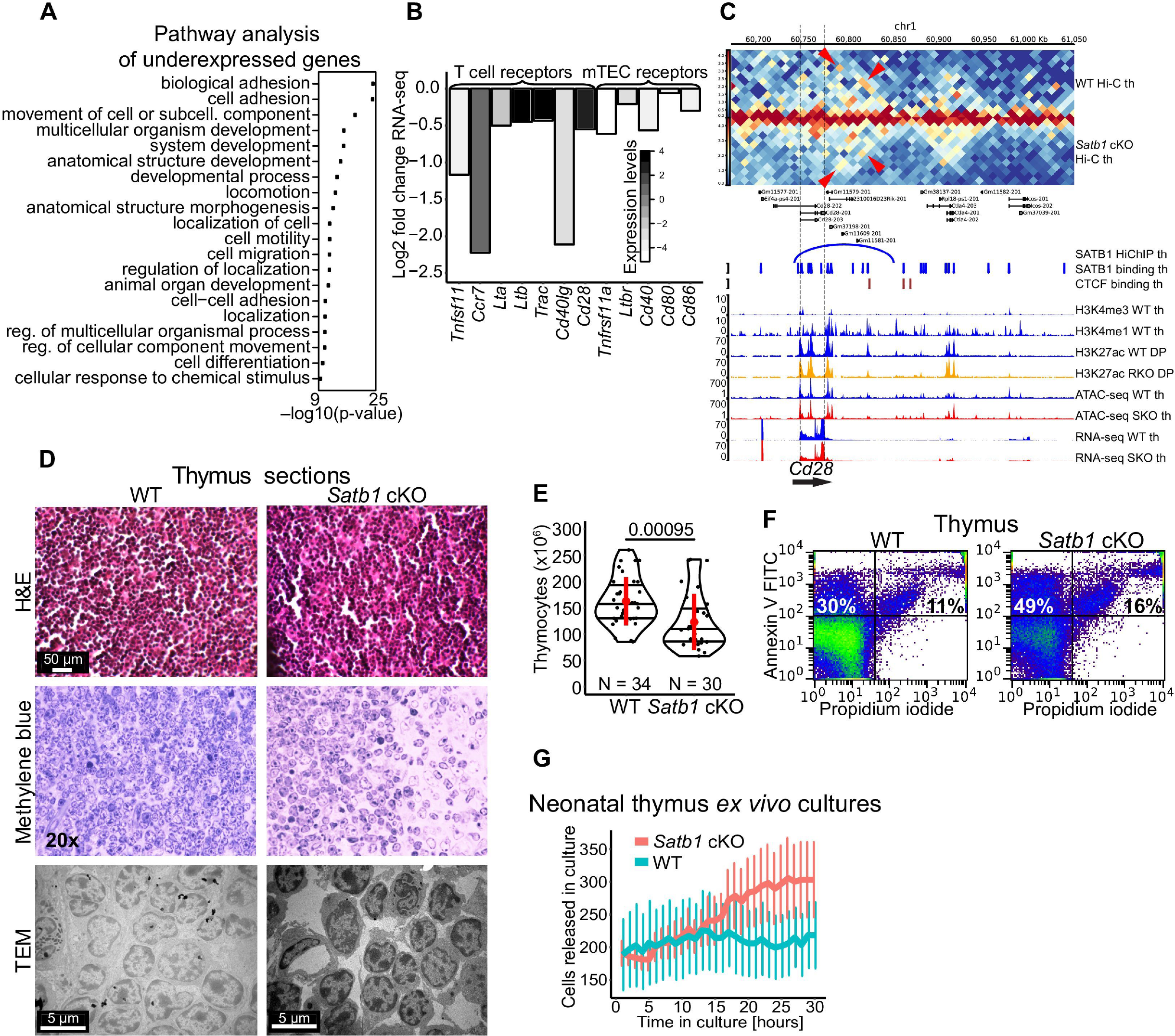
**Deregulated expression of genes encoding adhesion molecules and receptors in the Satb1 cKO thymocytes** (A) Pathway analysis for the genes differentially expressed in the *Satb1* cKO compared to WT thymocytes as deduced by RNA-seq. Biological adhesion was the top gene ontology pathway in *Satb1* cKO underexpressed genes. (B) Important thymic receptors were underexpressed in *Satb1* cKO. (C) *Cd28* gene regulated via SATB1-mediated promoter-enhancer chromatin loops. There was a SATB1-mediated regulatory loop and correspondingly increased interactions in the WT Hi-C which were decreased in the *Satb1* cKO Hi-C. Legend: th – thymocytes, DP – CD4^+^CD8^+^ T cells, RKO – *Rad21*^fl/fl^*Cd4-*Cre^+^ and SKO – *Satb1*^fl/fl^*Cd4-*Cre^+^. (D) Thymic sections displaying a disrupted thymic environment in the *Satb1* cKO. Thymic cryosections stained with Hematoxylin & Eosin (H&E) and with methylene blue indicated the decreased cellular density and decreased contacts between cells in the *Satb1* cKO compared to WT thymus. The transmission electron microscopy images are representative of four biological replicates to underscore the missing cellular contacts between the *Satb1* cKO cells. (E) Number of thymocytes in WT and *Satb1* cKO mice. The horizontal lines inside violins represent the 25^th^, 50^th^ and 75^th^ percentiles. The red circle represents the mean ± s.d. *P* values by Wilcoxon rank sum test. (F) The thymus of the *Satb1* cKO animals contained more apoptotic and necrotic cells as deduced by flow cytometry analysis using Annexin V and propidium iodide staining. (G) Neonatal thymi from WT and *Satb1* cKO mice were cultivated for 30 hours. An image was taken every hour to monitor the rate of T cell exit from the thymus. The error bars represent the standard error of the mean.

## Discussion

The adaptive immune response relies on the accurate developmental coordination of several alternative cell lineage fates. 3D genome organization in T cells represents a crucial denominator for this coordination (Spilianakis and Flavell, 2004; Spilianakis et al., 2005). In this work, we described the regulatory chromatin network of developing T cells and identified SATB1 protein being enriched at the anchors of regulatory chromatin loops. We performed a systematic genome-wide analysis of SATB1 roles in T cells. First, we compared the chromatin organization role of SATB1, to that of the conventional genome organizer CTCF. Utilizing a plethora of research approaches, we demonstrated that SATB1 establishes a finer-scale organizational layer, built upon the pre-existing scaffold mediated by other architectural proteins. Depletion of conventional genome organizers such as CTCF (Nora et al., 2017) or cohesin (Rao et al., 2017; Schwarzer et al., 2017) resulted in vast deregulation of TADs, however it did not show dramatic changes in gene expression as one would expect. On the contrary, SATB1 depletion did not result in any changes of TADs or high order chromatin organization, yet the long-range promoter-enhancer interactions were highly deregulated as well as the underlying transcriptional programs.

It was not clear so far whether SATB1 should be primarily assigned a role as an activator or a repressor. This characteristic makes SATB1 markedly similar to the ubiquitously expressed factor YY1. YY1 was also found enriched at promoters and enhancers, mediating their spatial contacts (Weintraub et al., 2017); however, depending on the cellular context it has also been found in association with Polycomb repressive complexes (Bracken and Helin, 2009). It would be interesting to further investigate the determinants of such functional ambiguity for such factors. In our experimental setup, we have demonstrated the activatory function of SATB1, in specifically mediating promoter-enhancer long-range chromatin interactions. Although the repressed nuclear environment of *Satb1* cKO cells was in agreement with those findings, we should note that in our study we were mostly focused on functions of the long SATB1 protein isoform. The presence of two SATB1 protein isoforms was recently described by our group (Zelenka et al., Submitted) and it could be another reason, aside from various post-translational SATB1 variants (Kumar et al., 2006; Zelenka and Spilianakis, 2020), supporting its functional ambiguity. We showed that the long SATB1 protein isoform had a higher propensity to undergo phase transitions compared to the short isoform (Zelenka et al., Submitted). Considering the proposed model of transcriptional regulation via liquid-liquid phase separated transcriptional condensates (Cho et al., 2018; Sabari et al., 2018), we reasoned that even the subtle differences in biophysical properties between the two SATB1 isoforms may play an important regulatory role. Following these observations, we hypothesize a model in which SATB1-mediated contacts between promoters and enhancers are attracted to the transcriptional condensates depending on the type and concentration of the SATB1 variants present. Post-translational modifications and/or different SATB1 isoforms would therefore regulate the whole process and under certain conditions, SATB1 could even function as a repressor. Thus, in comparison to other transcription factors capable of mediating long-range chromatin interactions, the SATB1’s mode of action may include one or more mechanisms that were proposed (Giammartino et al., 2020; Kim and Shendure, 2019; Stadhouders et al., 2019); i.e. via direct (Wang et al., 2012, 2014) or indirect (Fujii et al., 2003; Jangid et al., 2014; Kumar et al., 2005; Purbey et al., 2009; Yasui et al., 2002) oligomerization or via phase separation (Zelenka et al., Submitted). Overall, the presence of proteins like SATB1 with tissue-restricted expression profile, may represent the missing link between chromatin organization and tissue-specific transcriptional regulation.

One of our goals was to provide molecular mechanisms such as the 3D chromatin organization of T cells in order to explain the phenotypical malformations observed in the *Satb1* cKO mice. *Satb1*-deficient mice suffer from multiple health problems, markedly resembling autoimmunity. Previous research suggested a cell-extrinsic mechanism of autoimmunity based on the deregulation of regulatory T cells (Kitagawa et al., 2017). Here we presented that SATB1 is a regulator of several genes involved in T cell development, such as *Bcl6*, *Tcf7*, *Lef1*, *Cd6*, *Cd8*, *Lta*, *Ltb* and others. The individual deregulation of most of these genes would also result in deregulated immune responses. A great example for a cell-intrinsic mechanism of autoimmunity in the *Satb1* cKO is the SATB1-mediated spatial rearrangement of the TCRα enhancer and the TCR locus *per se*, controlling TCR recombination. A previous study linked the deregulation of the TCRα locus identified in the *Satb1* cKO to the downregulation of the *Rag1* and *Rag2* genes (Hao et al., 2015). In addition to this deregulation and the SATB1-mediated regulatory loops at the *Rag* locus, we also revealed the deregulation of chromatin accessibility and H3K27ac looping at the TCR locus. This was correlated with the SATB1-mediated chromatin loops and ultimately the defective usage of the different TCR segments. Since the recruitment of RAG proteins is based on the epigenetic status of a gene locus (Maman et al., 2016; Teng et al., 2015), we hypothesized that the altered TCR organization, due to missing SATB1-mediated loops, together with the disrupted recruitment of chromatin modifying complexes interacting with SATB1 (Fujii et al., 2003; Jangid et al., 2014; Kumar et al., 2005; Purbey et al., 2009; Yasui et al., 2002), would be critical contributors for the defective TCR arrangement in the *Satb1* cKO. Moreover, the impact of the 3D organization of TCR and BCR, necessary for proper VDJ recombination was recently highlighted (Peters, 2021; Rogers et al., 2021).

Apart from the TCRα locus, the most affected gene regarding the differential interaction analysis of regulatory loops between WT and *Satb1* cKO thymocytes was *Bcl6*. BCL6 represents a master regulator of Tfh (Johnston et al., 2009; Nurieva et al., 2009; Yu et al., 2009) and innate-like T cells (Gioulbasani et al., 2020). We have demonstrated that the Tfh program is deregulated in the *Satb1* cKO utilizing several research approaches. Moreover, the blockade at stage 0 (ST0) of iNKT development in *Satb1*-deficient mice was previously reported (Kakugawa et al., 2017), collectively suggesting a potential link between SATB1-mediated regulation of *Bcl6* and these developmental programs. BCL6 is known to function as an antagonist of factors specifying other cell lineage fates (Vinuesa et al., 2016; Wu et al., 2018), especially of PRDM1 and underlying Th17 lineage specification (Johnston et al., 2009). Indeed, the increased IL-17 response we observed in the cytokine milieu of *Satb1* cKO mouse sera (Figure 1D), suggests a favored Th17 specification due to downregulation of BCL6. However, since the depletion of SATB1 and its underlying regulome took place already during the intra-thymic development (*Satb1*^fl/fl^*Cd4*-Cre^+^), we hypothesize that the increased IL-17 cytokine levels were primarily due to potentially elevated γδT17 cells. Based on the transcriptomic analysis depicted in Figure 5A (RNA-seq, WT/*Satb1* cKO) we have detected the overexpression of the TCRδ locus and correspondingly also of the Vγ4^+^ and Vγ6^+^ chains which are expressed in γδT17 cells (Buus et al., 2016; Haas et al., 2012; Muñoz-Ruiz et al., 2017; Papotto et al., 2017). Moreover, the most overexpressed gene in *Satb1* cKO thymocytes, as indicated by our RNA-seq experiments, was *Maf* (encoding c-MAF; log2FC = 3.696 in female thymus and log2FC = 7.349 in male DP cells), which was shown to be essential for the commitment of γδT17 cells (Zuberbuehler et al., 2019).

Our findings point to the regulatory overlap between TCR recombination and transcriptional activity of master regulator genes in T cells, collectively orchestrated via spatial chromatin arrangements controlled by SATB1 and ultimately leading to the control of developmental decisions in the thymus. We provide a unique report on the functional intersection between CTCF and a tissue-specific genome organizer such as SATB1. Using the link between the altered 3D enhancer network and the physiological deregulation of *Satb1* cKO animals, we demonstrate the importance of the functional layer of chromatin organization provided by transcription factors such as SATB1. Our goal is to stir up a discussion about the existence of other tissue- or cell type-restricted factors, potentially contributing to the higher complexity and direct regulatory potential of the 3D chromatin architecture.

## Acknowledgements

We would like to thank Elena Deligianni for the neonatal thymi image acquisition, Sevasti Papadogiorgaki and George Chalepakis for transmission electron microscopy and George Garinis and Manouela Kapsetaki for fruitful discussions. Molecular graphics and analyses were performed with UCSF Chimera, developed by the Resource for Biocomputing, Visualization, and Informatics at the University of California, San Francisco, with support from NIH P41-GM103311. This work was supported by the European Union (European Social Fund ESF) and Greek national funds through the Operational Program ‘Education and Lifelong Learning’ of the National Strategic Reference Framework (NSRF) Research Funding Program ARISTEIA [MIRACLE 42], by FONDATION SANTE (X-COAT) and by Chromatin3D-H2020-MSCA-ITN (GA642934). D.P. was supported by the Polish National Science Centre (2019/35/O/ST6/02484 and 2020/37/B/NZ2/03757), Foundation for Polish Science co-financed by the European Union under the European Regional Development Fund (TEAM to DP), and by Warsaw University of Technology within the Excellence Initiative: Research University (IDUB) programme. The funders had no role in study design, data collection and analysis, decision to publish, or preparation of the manuscript.

## Author contributions

T.Z. and C.S. designed the study. T.Z. performed the genomics and immunofluorescence experiments. T.Z. and A.K. performed the computational analyses. S.F. consulted for library construction and performed sequencing experiments. P.T. and D.T. created the *Satb1* cKO mouse. T.Z. and D.T. performed animal, histology and flow cytometry experiments. D.A.P. performed the western blot experiments. I.R.T. performed computational 3D modeling experiments. D.P. and C.N. consulted the computational analyses. T.Z. wrote the original manuscript. C.S. supervised the work, obtained funding and corrected the manuscript. All authors read, discussed and approved the manuscript.

## Data availability

All genomics experiments are deposited in Gene Expression Omnibus database under accession number GSE173476. Other datasets will be provided upon a reasonable request.

## Methods

### Animals and isolation of thymocytes

All experiments were conducted in accordance with the Laboratory Animal Care and Ethics Committee of IMBB-FORTH. Animal work was approved by the IMBB Institutional Animal Care and Ethics Committee. All the experiments were performed on mice with C57BL/6 background. The generation of *Satb1*^fl/fl^ mice was previously described (Denaxa et al., 2012). The *Satb1* cKO (conditional knockout) mouse under study was created by crossing the *Satb1*^fl/fl^ mouse with a *Cd4-*Cre transgenic animal. The animals used for the experiments were 4-8 weeks old, unless otherwise specified. Primary thymocytes were resuspended by rubbing and passing the thymus through a 40 µm cell strainer (Falcon, 352340) into 1X PBS buffer. Cells were washed twice with 1X PBS: cells were centrifuged at 500 g, 4°C for 5 minutes, resuspended in 10 ml of 1X PBS and both steps were repeated. Prepared thymocytes were either used directly for experiments or fixed with 1% methanol-free formaldehyde (Pierce, 28908) at room temperature (RT) for 10 minutes while rocking. To quench the reaction, glycine was added to 0.125 M final concentration and incubated at RT for 5 minutes, while rocking. Cells were centrifuged at 1,000 g, 4°C for 5 minutes and washed twice with ice cold 1X PBS.

### Flow cytometry

#### Characterization of T cell populations in the *Satb1* cKO

Depending on the experiment, we used either thymocytes or splenocytes. Splenocytes were isolated in the same way as thymocytes, but they were further resuspended in plain water for 3 seconds to lyse erythrocytes, with immediate dilution by HBSS (Gibco, 14180) to a final 1X concentration. One million cell aliquots were distributed into 5 ml polystyrene tubes (BD Falcon 352052). For the experiments probing the percentage of apoptotic cells in tissues, we followed the PI/Annexin protocol (Biolegend, 640914). For staining with antibodies, we washed the cells once with Staining Buffer (1X PBS, 2% FBS, 0.1% NaN3) and then stained in 100 μl of Staining Buffer with 1 μl of antibodies at 4°C for 30 minutes. The stained cells were washed with excess of Wash Buffer (1X PBS, 0.5% FBS) and then analyzed on FACSCalibur flow cytometer. The antibodies used in flow cytometry experiments were anti-DNA PI-conjugated (Biolegend-79997), anti-Phosphatidylserine FITC conjugated (Biolegend-640906), anti-CD4 PE conjugated (Pharmingen-553730), anti-CD8a APC conjugated (Biolegend-100712), anti-CD44 PE conjugated (Pharmingen-553134) and anti-CD62L FITC conjugated (Biolegend-104406).

#### Infiltration of CD4^+^ cells in pancreas

Pancreas was isolated from three WT and three *Satb1* cKO mice of 120-136 days of age. Pancreas was cut in pieces and digested in 5 ml of 1 mg/ml collagenase (SIGMA, C2674) in PBS solution at 37°C for 30 minutes. Samples were washed twice with 5% FBS in PBS and filtered through a polypropylene mesh. After centrifugation, cell pellets were resuspended in 1 ml of 0.05% Trypsin solution and incubated for 5 minutes at 37°C. Cells were washed twice with ice-cold PBS and eventually filtered through a 40 µm cell strainer and blocked in 5 ml of 5% FBS in PBS for 30 minutes at 4°C. Cells were stained with 1:200 CD4-PE and CD8-APC for 30 minutes at 4°C and then washed twice with 0.5% FBS in 1X PBS. Lastly, cells were resuspended in 2% FBS in 1X PBS and analyzed by flow cytometry.

### Characterization of the cytokine milieu

Cytokines were characterized and quantified from serum of 16 female mice (5 WT, 11 *Satb1* cKO) of varying age 1-7 months by the LEGENDplex (13-plex) Mouse Th Cytokine Panel V02 (Biolegend, 740741; Lot B289245) according to the manufacturer’s instructions. Data were analyzed by the provided software LEGENDplex 8.0.

### Intraperitoneal glucose tolerance test

Groups of four WT and five *Satb1* cKO animals of 85 days (±8) of age were fasted for 6 hours. Weight and blood glucose levels were measured before and after the fasting period. 10% dextrose solution was injected intraperitoneally – the volumes were adjusted individually for each animal in order to inject 2 g of dextrose per kg of body mass. Blood (taken from tail) glucose levels were measured at given time points using Bayer Contour XT machine with Bayer Ascensia Contour Microfill Blood Glucose Test Strips. Animals were sacrificed and their pancreas was used for histology sections to demonstrate the disturbance of the islets of Langerhans.

### Histology and tissue sectioning

Samples were fixed in 4% formaldehyde in 1X PBS (pH 7.4) for 12-15 hours at 4°C. Tissues were rinsed in PBS and stored in PBS at 4°C until embedding. For embedding, samples were dehydrated for 30 minutes in 70% ethanol, 2x30 minutes in 90% ethanol and 3x30 minutes in 100% ethanol – all at RT while stirring. Specimens were cleared for 2x60 minutes in xylol and then impregnated for 1-2 hours at 58°C with paraffin. Samples were positioned in embedding moulds and left overnight to harden. Samples were sectioned on a sliding microtome to achieve 5-10 μm thin sections. The prepared sections on glass slides were deparaffinized for 30 minutes at 65°C and then for 2x 30 minutes in Neo-Clear (Merck Millipore, 109843). Samples were rehydrated for 2x 10 minutes in 100% ethanol, 1x 5 minutes in 90% ethanol, 1x 5 minutes in 70% ethanol, 1x 5 minutes in 50% ethanol, 1x 5 minutes in 30% ethanol and 1x 5 minutes in 1X PBS. Samples were immersed in Haematoxylin bath for 5 minutes in dark and then washed by running water for 5-10 minutes and in 1X PBS for 1 minute. Next, samples were immersed in Eosin bath for 30 seconds in dark and then washed by running water for 5-10 minutes and in 1X PBS for 1 minute. Samples were dehydrated again by dipping ten times in 30%, 50% and 70% ethanol and then incubated for 2x2 minutes in 100% ethanol. Eventually, samples were incubated for 2x10 minutes in xylene and then mounted using Entellan® new (Merck Millipore, 107961).

### Transmission electron microscopy

For scanning electron microscopy (SEM), fresh thymi were cut into small blocks. Briefly, tissue was fixed for 2 hours with 2% paraformaldehyde – 2% glutaraldehyde in 0.1 M sodium cacodylate buffer. Samples were post-fixed overnight in 1% osmium tetroxide (OTO method) and dehydrated in a graded series of ethanol. Specimens were coated in gold, mounted on aluminum stubs and examined with a JEOL JSM6390 LV scanning electron microscope (Peabody, MA) using an accelerating voltage of 15 kV.

### Detection of autoantibodies

The WT pancreas sample was prepared as previously described. 5 μm thick sections were deparaffinised at 55°C for 8 min and then processed in the following solutions: 2x 3 minutes in Neo-Clear, 2x 3 minutes in 100% ethanol, 1x 3 minutes in 95% ethanol, 1x 3 minutes in 70% ethanol, 1x 3 minutes in 50% ethanol. Samples were then rinsed with water and carefully dried with a paper towel. Tissue was circled with a PAP pen, let dry for 1 minute and then dipped in PBS for 2 minutes. Antigens were retrieved by incubation with 130 μl of TE-Triton-PK solution (2 ml TE buffer, 10 μl 0.5% Triton X-100, 40 μg Proteinase K) in a humidified chamber at 37°C, for 12 minutes. Samples were then washed twice with TBST buffer (10 mM Tris-HCl – pH 8.0, 150 mM NaCl, 0.05% Tween-20) for 3 minutes each. Samples were blocked by incubation with 5% normal goat serum (NGS) in TBST buffer at RT for 30 minutes in a humidified chamber. Samples were incubated at 4°C overnight with blood serum collected from two WT and four *Satb1* cKO animals of 4-7 months of age. Serum was diluted 1:10 in 5% NGS-TBST and 5% NGS-TBST was used as a negative control. Samples were washed twice with TBST, 5 minutes each. Samples were incubated with a goat anti-mouse IgG antibody (H+L; Invitrogen, A-11032) conjugated with Alexa Fluor 594, diluted 1:500 in TBST at RT for 1 hour. Samples were washed three times, 5 minutes each, with TBST and incubated with 1 μM DAPI solution in 5% NGS-TBST at RT for 10 minutes. Samples were washed three times, 5 minutes each, with TBST and mounted with Mowiol on glass slides.

### Hi-C and HiChIP experiments

#### Generation of proximity-ligated contacts

A biological duplicate was used for each sample. Both Hi-C and HiChIP experiments were performed identically until the chromatin immunoprecipitation step. Aliquots of 10 million isolated thymocytes resuspended in 1X PBS were fixed by adding 1/10^th^ volume of fixation butter [11% methanol-free formaldehyde (Pierce, 28908), 100 mM NaCl, 1 mM EDTA, 0.5 mM EGTA, 50 mM Hepes pH 8.0] with rocking at RT for 10 minutes. To quench the reaction, glycine was added to 0.125 M final concentration and incubated at RT for 5 minutes, while rocking. After two washes with 1X PBS, cell pellet was resuspended in 500 μl of ice-cold Hi-C Lysis Buffer (10 mM Tris-HCl – pH 8, 10 mM NaCl, 0.2% NP40, 0.5 mM PMSF) and rotated at 4°C for 1.5 hours. Cells were centrifuged at 2,500 g, at 4°C for 5 minutes and the supernatant was discarded. The cell pellet was washed once with 500 μl of ice-cold Hi-C Lysis Buffer and then resuspended in 100 μl of 0.5% SDS. Cells were incubated at 62 °C for 10 minutes and then combined with 296 μl of H2O and 50 μl of 20% Triton X-100. Samples were incubated at 37°C for 15 minutes and then combined with 50 μl of 10X DpnII Buffer and 200 U of DpnII restriction enzyme (NEB, R0543M) and digested for additional 16 hours at 37 °C while shaking (160 rpm). The restriction enzyme was inactivated at 62°C for 20 minutes and the nuclei were centrifuged at 2,500 g, at 4 °C for 6 minutes. The supernatant was discarded and the nuclei were resuspended in 300 μl Fill-in Buffer containing 30 μl Klenow Buffer 10X (NEB, M0210L), 15 μl1 mM Biotin-16-dCTP (Jena Bioscience, NU-809-BIO16-L), 1.5 μl 10 mM dATP (Promega, U1240), 1.5 μl 10 mM dGTP (Promega, U1240), 1.5 μl 10 mM dTTP (Promega, U1240), 12 μl 5 U/μl DNA Polymerase I, Klenow Fragment (NEB, M0210L) and 238.5 μl water. The biotinylation mixture was incubated at 37°C for 30 minutes with rotation. SDS was added to a final concentration of 0.5% to inactivate the Klenow enzyme. Triton X-100 was added to 1% final concentration and samples were incubated at 37°C for 5 minutes. Samples were centrifuged at 2,500 g, at 4 °C for 10 minutes and the supernatant was discarded. The nuclei pellet was resuspended in the Ligation Buffer containing 120 μl 10X NEB T4 DNA Ligase Buffer supplemented with 10 mM ATP (NEB, B0202), 60 μl 20% Triton X-100 (1% final), 6 μl 2% (20 mg/ml) BSA, 40 μl 30% PEG 6,000 (1% final), 5 μl 400 U/μl T4 DNA Ligase (NEB, M0202L) and 969 μl water. The samples were incubated for 6 hours at RT with mild rotation. Nuclei were centrifuged at 2,400 g, at RT for 15 minutes and the supernatant was discarded. The pellet was resuspended in 60 μl Lysis Buffer (1% SDS, 50 mM Tris-HCl – pH 8, 20 mM EDTA, 1X protease inhibitors) and incubated at RT for 15 minutes. The lysate was diluted to 600 μl using ice cold TE Buffer supplemented with protease inhibitors and then sonicated with a Labsonic M – Tip sonicator for 3 cycles (30 seconds ON/OFF, 40% power). The sonicated material was centrifuged at 16,000 g, at RT for 15 minutes and the supernatant was collected into a new tube. Samples from the same genotype were merged and then split again: separately 100 μl for Hi-C and 450 μl for HiChIP.

**Hi-C** samples were combined with two volumes of Hi-C Elution Buffer (10 mM Tris-HCl – pH 8, 5 mM EDTA, 300 mM NaCl, 1% SDS) and incubated at 65°C overnight. Decrosslinked material was diluted to 500 μl with TE Buffer and treated with RNase A and Proteinase K as previously described. DNA was purified using a ChIP DNA Clean & Concentrator kit following the manufacturer’s instructions (Zymo Research, D5205). Purified DNA was quantified using Qubit dsDNA BS Assay Kit (Invitrogen, Q32853) checked on an agarose gel for shearing efficiency and 100 μg were used for library construction.

**HiChIP** samples were combined with Triton X-100 to 1% final concentration and samples were incubated at 37°C for 15 minutes. Samples were combined with an equal volume of 2X ChIP Binding Buffer (20 mM Tris-HCl – pH 8, 2 mM EDTA, 0.2 % sodium deoxycholate, 2X protease inhibitors). Chromatin preclearing and antibody binding to magnetic beads were performed as described in the ChIP protocol. The following antibodies were used for the immunoprecipitation step: 8 μg of custom-made David’s Biotechnologies SATB1 long isoform, 7 μg Abcam (ab70303) anti-CTCF and 2 μg Abcam (ab4729) anti-H3K27ac.

Antibody-coupled beads were incubated with chromatin at 4°C with rotation for 16 hours. Beads were washed five times with ice cold RIPA Buffer (50 mM Hepes pH 8, 1% NP-40, 0.70% Na-Deoxycholate, 0.5 M LiCl, 1 mM EDTA, protease inhibitors) and then twice with TE Buffer. After the first wash with TE Buffer, resuspended beads were transferred into a new tube. Immune complexes bound to beads were eluted in 125 μl Hi-C Elution Buffer (10 mM Tris-HCl – pH 8, 5 mM EDTA, 300 mM NaCl, 1% SDS) at 65°C for 16 hours. Decrosslinked material was diluted to 250 μl with TE Buffer and treated with RNase A and Proteinase K as previously described. DNA was purified using a ChIP DNA Clean & Concentrator kit following the manufacturer’s instructions (Zymo Research, D5205). Purified DNA was quantified using Qubit dsDNA HS Assay Kit (Invitrogen, Q32854) and used for library construction.

#### Biotin pull-down and library construction

Samples were brought to 25 μl with water. 5 μl and 20 μl for HiChIP / Hi-C samples, respectively, of Dynabeads MyOne Streptavidin C1 beads (Invitrogen, 65001) were washed with 500 μl Tween Wash Buffer (5 mM Tris-HCl – pH 7.5, 0.5 mM EDTA, 1 M NaCl, 0.05% Tween-20). Beads were resuspended in 25 μl of 2X Biotin Binding Buffer (10 mM Tris-HCl – pH 7.5, 1 mM EDTA, 2M NaCl) and combined with samples. Samples were incubated at RT for 20 minutes. Beads were separated on a magnet and washed twice with 400 μl of Tween Wash Buffer. Beads were washed by 100 μl of 1X TD Buffer (10 mM Tris-HCl – pH 7.5, 5 mM MgCl_2_, 10% Dimethylformamide). Beads were resuspended in 25 μl of 2X TD Buffer and combined with Tn5 enzyme from the Nextera DNA Sample Preparation Kit (Illumina, FC-121-1030) and water to final volume 50 μl. The amount of Tn5 enzyme was adjusted according to the input DNA amount: 4.5 μl for Hi-C libraries, 1.5 μl for SATB1 HiChIP and 1 μl for other HiChIP libraries. The reaction was incubated at 55°C for 10 minutes. The beads were collected with a magnet and the supernatant was discarded. Beads were resuspended in 300 μl of Strip Buffer (0.15% SDS, 10 mM Tris-HCl – pH 8, 50 mM EDTA) and incubated for 5 minutes at RT to strip off and deactivate Tn5. Beads were washed once with 400 μl of Tween Wash Buffer and once with 500 μl of 10 mM Tris-HCl (pH 8.0). The beads were resuspended in 50 μl of the following PCR master mix with indexed primers from the Nextera DNA Sample Preparation Index Kit (Illumina, FC-121-1011): Phusion HF 2X (NEB, M0531L) 25 μl, Nextera Index 1 (N7XX 5.5 μM) 1 μl (1.5 for Hi-C), Nextera Index 2 (N5XX 5.5 μM) 1 μl (1.5 for Hi-C) and water 23 μl (22 for Hi-C). The PCR reaction was performed following the program 72°C for 5 minutes and repeated cycles of 98°C for 15 seconds, 63 °C for 35 seconds, 72 °C for 1 minute. The number of PCR cycles was estimated based on post-ChIP quantification and amplification was 6 cycles for Hi-C libraries, 11 cycles for SATB1 HiChIP and 13 cycles for the other HiChIP libraries. DNA libraries were purified and size-selected using AMPure XP beads, quantified by Qubit and analyzed on a Bioanalyzer, as previously described. The DNA Libraries were sequenced on an Illumina® HiSeq 4000 2x 75 bp platform by the sequencing facility at IKMB, Kiel University, Germany.

#### Data processing

Raw reads were mapped with bowtie2 (Langmead and Salzberg, 2012) to the mm10 genome and fully processed using the HiC-Pro pipeline (version 2.11.1; Servant et al., 2015) with default parameters. All biological replicates were processed individually to assess their quality and then combined for downstream analyses and visualization. The HiChIP datasets were additionally processed by FitHiChIP (Bhattacharyya et al., 2019). Unless stated otherwise, the following parameters were used to call HiChIP loops: 5,000 kbp resolution, 20000-2000000 distance threshold, FDR 0.01, coverage specific bias correction, merged nearby peak to all interactions. Differential interacting areas between SATB1 and CTCF HiChIP matrices at 100 kbp and 500 kbp resolution were analyzed using diffHic (Lun and Smyth, 2015).

For the differential analysis of H3K27ac WT and *Satb1* cKO HiChIP datasets, we utilized the differential analysis pipeline from FitHiChIP (Bhattacharyya et al., 2019) and only utilized loops that were classified as differential in 3D but not in 1D. These were loops that showed differences in interaction counts but no significant differences in H3K27ac occupancy at loop anchors. Since some regions contained H3K27ac loops from both under- and over-interacting categories, we furthermore calculated a difference between the number of under- and overinteracting loops.

Binding site datasets, needed to call HiChIP loops, were either derived from HiChIP data or an external ChIP-seq dataset. The SATB1 binding sites were extracted from another HiChIP experiment with >60% Dangling End Pairs (∼280 million reads), using the PeakInferHiChIP.sh script from FitHiChIP (--nomodel --extsize 147; filter peaks with >2.5 enrichment). SATB1 binding sites were compared to a published SATB1 ChIP-seq dataset (GSM1617950; Hao et al., 2015). Both our biological replicates and the published dataset revealed similar overrepresented categories in genome and gene ontology functional analyses as well as a high overlap of peaks and differentially expressed genes. The H3K27ac peaks were derived similarly from the HiChIP datasets (separately for two biological replicates and then merged). The CTCF peaks were also derived the same way and motif analysis with MEME (Bailey and Elkan, 1994) validated the high enrichment of the CTCF binding motif in the HiChIP derived peaks, confirming its specificity. However, due to a relatively low number of HiChIP-derived CTCF binding sites, for FitHiChIP loop calling and other computational analyses, we employed a CTCF ChIP-seq dataset from the ENCODE project (ENCFF714WDP; Dunham et al., 2012). Only for the visualization purposes, we used combined datasets (mergeBed command of bedtools; Quinlan and Hall, 2010) of our HiChIP-derived binding sites and publicly available ChIP-seq datasets for both SATB1 (GSM1617950; Hao et al., 2015) and CTCF (ENCFF714WDP; Dunham et al., 2012). Since our antibodies were specific for the long SATB1 isoform, we used the dataset combined with the public ChIP-seq to ensure that all types of SATB1 peaks were shown.

At the TCR locus, whole DNA segments are missing in some cells due to V(D)J recombination. Thus, in the cell population some regions of TCR are underrepresented compared to other genomic loci and this fact penalizes loop or binding site calling at the TCR locus. For this reason, for the TCR analysis we used different datasets with adjusted and more relaxed parameters to compensate for this. For the purpose of loop calling in HiChIP experiments, we utilized the combined binding site datasets of our HiChIP-derived binding sites and publicly available ChIP-seq datasets for both SATB1 (GSM1617950; Hao et al., 2015) and CTCF (ENCFF714WDP; Dunham et al., 2012). Furthermore, the following parameters were used: 10,000 kbp resolution, 20000-2000000 distance threshold, FDR 0.05, coverage specific bias correction, merged nearby peak to all interactions.

For operations with matrices, the datasets were processed using hicexplorer (Ramírez et al., 2018). Hi-C and HiChIP matrices were normalized to the smallest dataset from each compared pair, i.e. WT vs SKO Hi-C, SATB1 vs CTCF HiChIP and WT vs SKO H3K27ac HiChIP. Normalized matrices were corrected using *hicCorrectMatrix* based on the diagnostic plots applying a KR balancing method (Knight and Ruiz, 2013). The analysis of A/B compartments was done according to the original protocol (Lieberman-Aiden et al., 2009) using hicexplorer (Ramírez et al., 2018) and/or by HOMER (Heinz et al., 2010). Differential analysis of TADs was performed using TADCompare (Cresswell and Dozmorov, 2020) at 100 kbp resolution on matrices combined for both biological replicates to compare WT and *Satb1* cKO. As a control we also compared TADs called from individual replicates of each genotype. Visualization of matrices was done by hicexplorer (Ramírez et al., 2018), pyGenomeTracks and/or by Juicebox (Durand et al., 2016). APA scores (Rao et al., 2014) were calculated and visualized by SIPMeta (Rowley et al., 2020) and/or by Juicer Tools (Durand et al., 2016).

### Stranded-total-RNA sequencing

#### Experimental protocol

Freshly isolated thymocytes from female animals were resuspended in 1 ml of TRIzol Reagent (Invitrogen, 15596026) and RNA was isolated according to manufacturer’s protocol. The aqueous phase with RNA was transferred into a tube and combined with 10 μg of Linear Acrylamide (Ambion, AM9520), 1/10 of sample volume of 3M CH3COONa (pH 5.2), 2.5 volumes of 100% Ethanol and tubes were mixed by flipping. Samples were incubated at –80°C for 40 minutes. Samples were centrifuged at 16,000 g, at 4°C for 30 minutes. The supernatant was removed and the pellet was washed twice with 75% Ethanol. The air-dried pellets were resuspended in 40 μl RNase-free water and incubated at 55°C for 15 minutes to dissolve RNA. To remove any residual DNA contamination, RNase-free DNase Buffer was added to the samples until 1X final concentration together with 20 units of DNase I (NEB, M0303L) and incubated at 37°C for 20 minutes. Samples were then purified using RNeasy Mini Kit (Qiagen, 74104) according to the manufacture’s protocol. RNA quality was evaluated using Agilent 2100 Bioanalyzer with Agilent RNA 6000 Nano Kit (Agilent Technologies, 5067-1511). Libraries were prepared using an Illumina® TruSeq® Stranded Total RNA kit with ribosomal depletion by Ribo-Zero Gold solution from Illumina® according to the manufacturer’s protocol and sequenced on an Illumina® HiSeq 4000 (2x 75 bp).

#### Data processing

Raw reads were mapped to the mm10 mouse genome using HISAT2 (Kim et al., 2019). Only mapped, paired reads with a map quality >20 were retained. Transcripts were assembled with StringTie (Pertea et al., 2015) using an evidence-based Ensembl-Havana annotation file. Transcripts and genes were summarized using featureCounts (Liao et al., 2014) and statistically evaluated for differential expression using DESeq2 (Love et al., 2014). When application required an intra-sample transcript comparison, DESeq2 values were further normalized to the gene length. The functional analyses were performed by g:Profiler (Reimand et al., 2007). The plots depicting enriched BP terms and KEGG pathways were generated by presenting the top 20 pathways/terms with the lowest p-values.

### ATAC-seq

#### Experimental protocol

A biological triplicate was used for each genotype. The ATAC-seq experiment was performed according to the Omni-ATAC protocol previously published (Corces et al., 2017), with modifications. Murine thymocytes were isolated as previously described, without fixation. To ensure the presence of only viable cells, cells were separated using Lympholyte®-M (Cedarlane, CL5030) according to the manufacturer’s protocol. Isolated cells were washed twice with 1X PBS and aliquots of 10,000 cells were used for analysis. The cell pellet was gently resuspended by pipetting up and down three times in 50 µl of ice cold ATAC-RSB-NTD Buffer (10 mM Tris-HCl – pH 7.5, 10 mM NaCl, 3 mM MgCl_2_, 0.1% NP40, 0.1% Tween-20, 0.01% Digitonin) and incubated on ice for 3 minutes. Cell lysis was stopped by adding 1 ml of cold ATAC-RSB-T Buffer (10 mM Tris-HCl – pH 7.5, 10 mM NaCl, 3 mM MgCl_2_, 0.1% Tween-20) and inverting the tube three times to mix. Nuclei were centrifuged at 1,000 g, at 4°C for 10 minutes. The pellet was resuspended in 50 µl of Transposition Mix [25 µl 2X TD buffer (20 mM Tris-HCl – pH 7.6, 10 mM MgCl2, 20% Dimethyl Formamide – before adding DMF, the pH was adjusted to 7.6 with 100% acetic acid), 2.5 µl transposase (100 nM final), 16.5 µl PBS, 0.5 µl 1% digitonin, 1 µl 5% Tween-20, 4.5 µl H2O] by pipetting up and down six times. The reaction was incubated at 37°C for 30 minutes with occasional pipetting. DNA was purified with a DNA Clean & Concentrator-5 Kit (Zymo Research, D4013) according to the manufacturer’s protocol. DNA was eluted in 20 µl of Elution Buffer and all the material was used in a PCR reaction, together with 25 µl Phusion HF 2X Master Mix (NEB, M0531L) and 2.5 µl of each Nextera Index 1 (N7XX) and Nextera Index 2 (N5XX) primers from a Nextera DNA Sample Preparation Index Kit (Illumina, FC-121-1011). PCR was performed according to the following protocol: 72°C for 5 minutes, 98°C for 1 minute and 5 cycles of 98 °C for 15 seconds, 63 °C for 35 seconds, 72 °C for 1 minute. The samples were put on ice and 5 µl of the pre-amplified mixture was combined with 15 µl of qPCR Master Mix using the following set-up: water 3.25 μl, primer ad1 0.5 μl, primer ad2 0.5 μl, 20x SYBR Green 0.75 μl, Phusion HF 2x Master mix 5 μl and pre-amplified sample 5 μl. The qPCR reaction was run for 20 additional cycles following the program 98°C for 1 minute and 5 cycles of 98°C for 15 seconds, 63 °C for 35 seconds, 72 °C for 1 minute.

Based on the Rn (Fluorescence) vs Cycle linear plot, a cycle with 1/4 up to 1/3 of the maximum fluorescence level was determined. This was the number of additional PCR cycles to run on the pre-amplified libraries which were stored on ice until this point. The final amplified libraries were purified using a DNA Clean & Concentrator-5 Kit (Zymo Research, D4013) according to the manufacturer’s protocol and eluted in 20 µl of Elution Buffer. Libraries were quantified by Qubit and analyzed on a Bioanalyzer as previously described, followed by two-sided size selection using AMPure Beads. Libraries were sequenced on an Illumina® HiSeq 4000 2x 75 bp platform by the sequencing facility at IKMB, Kiel University, Germany.

#### Data processing

To acquire BAM files and ATAC-seq peaks, raw data were fully processed by the esATAC pipeline (Wei et al., 2018). To identify the regions with differential accessibility between genotypes, all esATAC called peaks across samples and replicates were pooled and tested for differences in accessibility levels. ATAC-seq counts were calculated for each peak, each replicate and each condition using FeatureCounts (Liao et al., 2014) and used as an input for edgeR (Robinson et al., 2010) with the standard parameters. A cutoff of |logFC|≥1 and p-value ≤0.01 was used to determine the differentially accessible regions.

To assess the accessibility around SATB1 binding sites BigWig files were generated. Mitochondrial reads and PCR duplicates were removed from all bam files using samtools (Li et al., 2009) and Picard MarkDuplicates (http://broadinstitute.github.io/picard/), respectively. Low quality and unmapped reads were removed using the following samtools command: samtools view h -b -F 1804 -f 2 -q 30. The final bam files were merged for the two conditions using samtools and RPKM normalized BigWig files were generated using deeptools (Ramírez et al., 2016) with the following parameters: -of bigwig --effectiveGenomeSize 2652783500 --normalizeUsing RPKM -bl New_merged_mm10.blacklist.bed -bs 1. mm10 blacklisted regions were downloaded from ENCODE. Moreover, using the merged bam files, BigWig files with the log2 ratio between the normalized reads of *Satb1* cKO and WT thymocytes were generated using deeptools bamCompare. The parameters were the same as above. SATB1 binding sites were first centered and then extended by 250 bp upstream and downstream. For each bp position, ATAC-seq signal was calculated using the generated BigWig files. Similarly, each TSS of each gene was centered and extended by 1 kbp upstream and downstream. For each base of each gene, an average log2 fold change of normalized accessibility score (*Satb1* cKO vs WT) was plotted. To calculate the accessibility changes along the entire genes, genes and upstream and downstream regions were divided into bins and average log2 fold change of normalized accessibility score for each bin was plotted.

### Immunofluorescence experiments

Glass coverslips were coated by dipping in 0.1 mg/ml poly-D-lysine solution (Sigma Aldrich, P6407). Freshly isolated thymocytes were attached to the coated coverslips. Attached cells were washed once with 1X PBS and then fixed for 10 minutes on ice with 4% formaldehyde (Pierce, 28908) in 1X PBS. Fixed cells were permeabilized with 0.5% Triton-X in 1X PBS for 5 minutes on ice. Cells were washed three times with 1X PBS for 5 minutes each and blocked for 30 minutes at RT with Blocking Buffer [0.4% acetylated BSA (Ambion, AM2614) in 4X SSC] in a humidified chamber. Cells were incubated for 1.5 hours at RT with an antibody against Hp1α (Merck Millipore, MAB3584, 1:500 dilution) and RNA Polymerase II (Santa Cruz, sc-900, 1:50) in Detection Buffer (0.1% acetylated BSA, 4X SSC, 0.1% Tween 20) in a humidified chamber. The excess of antibodies was washed away by three washes for 5 minutes each with Washing Buffer (4X SSC, 0.1% Tween 20). Cells were incubated for 60 minutes at RT with a goat anti-rabbit antibody conjugated with Alexa Fluor 488 (1:250) and a goat anti-mouse antibody conjugated with Alexa Fluor 647 (1:250) in Detection Buffer (0.1% acetylated BSA, 4X SSC, 0.1% Tween 20) in a humidified chamber. The excess was washed away by three washes for 5 minutes each with Washing Buffer (4X SSC, 0.1% Tween 20). The cells were mounted in a hardening ProLong Gold medium with DAPI (Invitrogen, P36935). Images were taken using an inverted microscope DMI6000 CS with laser scanning confocal head Leica TCS SP8, equipped with a 63x/1.40 oil immersion objective. Images were analyzed using the Fiji software (Schindelin et al., 2012). Cells were manually selected and signal was measured as an integrated signal density from summed z-stacks.

### Cultivation of neonatal thymi

Thymi of neonatal WT and *Satb1* cKO mice were collected and embedded in collagen. Thymi were cultivated in a medium (10% FBS, RPMI, Hepes, Pen-Strep, Glutamine, β-mercaptoethanol) for 30 hours and monitored with the Operetta high content screening microscope (PerkinElmer) to detect cells exiting the thymus. Images were pre-processed with the in-built analysis software Harmony 4.1 and then analyzed with custom-made macros in Fiji software (Schindelin et al., 2012). Random shifts were corrected by a StackReg plugin (Rigid Body transformation). Areas without any distortion were selected and cells outside the thymus were counted using Find Maxima function of Fiji. Each selection was normalized to the area size and for each animal the selections were averaged. The final result represents an average from different animals for each genotype.

### Linear regression model

A linear regression model was built in R and used to identify the impact of individual variables from our datasets on log2FC RNA-seq values. The predictors used are described below: Each gene was binned into 3 bins. Moreover, two extra bins upstream of the TSS of each gene (upstream region: –4 kbp to –2 kbp and promoter region: -2kb to TSS) and two extra bins downstream of the transcription termination site (TTS to +2 kbp and +2 kbp to +4 kbp) were used. SATB1 binding occupancy in WT cells was determined via a binary score for each bin of each gene. “1” indicated the presence of a SATB1 peak and “0” the absence of a SATB1 peak overlapping with the corresponding bin.

To quantify ATAC-seq signal differences between the *Satb1* cKO and WT thymocyte samples, the following calculation for each bin of each gene was used: log10[(*Satb1* cKO Normalized Reads + 0.01) / (WT Normalized Reads + 0.01)] * log2(Total Normalized Reads + 1)

To utilize SATB1 and CTCF loops as predictors, we assigned to each gene the number of times it overlapped with the anchors of a SATB1 or CTCF loop. In case that both anchors of the same SATB1/CTCF loop overlapped a gene, only one was counted. The number of times, the anchors of a SATB1 loop were found to connect an enhancer with a gene, was also used as a predictor. The same metrics were calculated for overinteracting and underinteracting H3K27ac loops.

The quality plots of the model are depicted in Figure S2A-D and the adjusted R-square of the model was 0.1128. The change of the Akaike Information Criterion (AIC, Figure S2E) estimated how the quality of the model was affected when all the predictors were kept intact except one. The y-axis corresponds to the removed predictor, while the x-axis indicates how the AIC for the new model was altered. An increase indicated that the predictor was “useful” for the model. Based on the AIC plot, neither SATB1 binding upstream and downstream of genes nor CTCF-mediated loops contributed to the accuracy of the model. On the other hand, differences in chromatin accessibility and connectivity via H3K27ac loops along with SATB1-mediated loops performed very well as predictors. Non-useful predictors were not used in the final model. The final model coefficients for the important predictors are displayed in Figure S2F. The sign of each coefficient indicates whether a predictor is associated with decreased (negative) or increased (positive) RNA levels in *Satb1* cKO. Genes present at anchors of overinteracting H3K27ac chromatin loops and/or with increased chromatin accessibility were associated with increased RNA levels in *Satb1* cKO. In contrast, the genes present at anchors of underinteracting H3K27ac chromatin loops or SATB1-mediated loops and/or genes bound by SATB1 were associated with reduced RNA levels.

### Additional bioinformatics analyses

#### Identification of SATB1 binding in H3K27ac HiChIP anchors

Loop anchors from WT H3K27ac HiChIP loops were pre-processed by extracting the anchors from both sides of loops and then they were merged into non-overlapping unique regions (mergeBed command of bedtools; Quinlan and Hall, 2010). The resulting regions were converted from mm10 to mm9 using CrossMap (Zhao et al., 2014) and were analyzed using the enrichment analysis ChIP-Atlas (Oki et al., 2018) against all the available murine ChIP-seq datasets from the blood cell type class and compared to 100X random permutations.

#### Publicly available ChIP-seq datasets

We utilized the CTCF ChIP-seq dataset from the ENCODE project (ENCFF714WDP; Dunham et al., 2012) and the SATB1 ChIP-seq (GSM1617950; Hao et al., 2015). For the analysis of enhancers, in relation to SATB1 and CTCF loops, we utilized the thymus-specific list of enhancers previously generated within the ENCODE project (GSE29184; Shen et al., 2012) – after extending the center of each enhancer by 50 bp upstream and downstream. Moreover, for visualization purposes we also utilized the H3K4me3 (ENCFF200ISF) and H3K4me1 (ENCFF085AXD) ChIP-seq datasets for thymus, from the ENCODE project (Davis et al., 2018). The H3K27ac ChIP-seq datasets were from WT and *Rad21*^fl/fl^*Cd4*-Cre^+^ DP cells (Ing-Simmons et al., 2015) and they were first converted from mm9 to mm10 using CrossMap (Zhao et al., 2014). An average file based on two biological replicates (GSM1504384+5 for WT and GSM1504386+7 for *Rad21* cKO) was created and used for visualization.

#### Overlap score calculation for HiCHIP loops

The overlap score between SATB1 and CTCF loops was calculated as (number of overlapping bp) / (bp size of a loop). Overlaps between the two types of loops were found using bedtools (Quinlan and Hall, 2010) and the above score was calculated using R. A score of one indicates either 100% overlap or engulfment of a loop mediated by one factor in a loop mediated by another factor. A score of zero indicates no overlap. In cases where a loop mediated by one factor intersected with multiple loops mediated by the other factor, the maximum score was used.

#### Functional analyses of gene lists

All gene ontology pathway analyses were performed with the R package gProfileR (Reimand et al., 2007). Twenty biological processes pathways with the lowest p-values were plotted.

#### Nucleosome binding

Nucleosome positions and factor occupancy was analyzed with NucleoATAC (Schep et al., 2015). NucleoATAC was run with standard parameters on merged ATAC-seq bam files, separately for each genotype. SATB1 and CTCF binding sites were used as the input for the occupancy analysis. Peaks were centered and extended 250 bp upstream and downstream prior to the analysis.

#### Transcriptional insulation scores

We investigated whether the expression of genes inside SATB1 and CTCF loops was different from the neighboring genes outside the loops. To test this, we established an insulation score by calculating the difference between the mean expression of genes inside loops [log10(counts / gene length + 0.01)] and the mean expression of genes found in the same size regions upstream and downstream from each loop. To compare the isolated values against a null distribution, loop coordinates were randomly shuffled across the genome and differences between the expression of genes residing inside the randomized loops with their neighbors were calculated as described above. If a shuffled loop was not containing any genes it was re-shuffled.

#### Permutation analyses

In order to construct a null statistical model (e.g. determination of common genes between two gene subsets, estimation of expected peak overlaps), permutation analyses were performed. In cases of overlaps between two files with genomic coordinates, the coordinates of one file were shuffled 1,000 times with the bedtools shuffle command (Quinlan and Hall, 2010). The new overlap for each iteration was calculated. The mean overlap count was used to determine enrichment. Finally, a p-value was calculated as follows: p-value = (X times the permutation overlap was higher than the actual overlap / 1000). The same was applied for overlaps between gene lists, with the exception of drawing out random genes instead of shuffling coordinates.

To generally test a null hypothesis for randomly selected values X and Y from two populations, the probability of X being greater than Y is equal to the probability of Y being greater than X, we used a nonparametric Wilcoxon rank sum test (ggsignif R package).

To statistically evaluate accessibility of SATB1 peaks (Figure 3B), each SATB1 HiChIP peak was centered and extended 250 bp upstream and downstream. An average read-depth normalized ATAC-seq score was calculated for each centered peak using the UCSC executable, bigWigSummary. Scores were log10 transformed. The process was repeated 100 times after random shuffling of the centered and extended peaks across the mm10 genome. The shuffleBed command from the bedtools suite (Quinlan and Hall, 2010) was used to shuffle the peaks. The average accessibility scores were calculated for the permuted and the "real" value distributions. A bootstrap p-value was calculated as: Number of randomly permuted mean values bigger than the "real" mean value / 100. In order to identify SATB1 peaks that occupied regions with reduced accessibility levels (Figure S1K), the aforementioned analysis was performed with 10 permutations. The average log10 transformed read-normalized accessibility scores of the ten random permutations was used as a cutoff for picking SATB1 peaks with low accessibility levels.

#### 3D computational modeling

The WT and *Satb1* cKO thymocyte Hi-C contact matrices, binned at a resolution of 20 kbp, were pre-processed by vanilla coverage normalization (Rao et al., 2014). An area delimited by two TADs (called by hicexplorer; Ramírez et al., 2018) at 150 kbp resolution and spanning chr16: 21420000-25340000) encompassing the *Bcl6* locus (chr16: 23965052-23988612) was used for 3D modeling of chromatin interactions, using TADbit Python library (Serra et al., 2017). Each contact matrix was modeled as a coarse-grained “beads-on-a-string” polymer model at an equilibrium scale of 0.01 nm/bp. First, we identified the optimal parameters to establish Z-score thresholds for attractive (*upfreq*) and repulsive (*lowfreq*) restraints, maximum inter-locus distance (*maxdist*) and maximum model physical distance, below which two model loci could be considered in contact with each other. Out of 500 models in total, we selected 100 models with the lowest amount of violated contact restraints. Next, we generated an ensemble based on the selected 100 models. Parameters were selected using a grid search approach aimed to optimize the correlation between the experimental input and the ensemble, as a model-derived contact map. The selected parameters were for WT: upfreq –0.3, lowfreq –1.0, maxdist: 550, cutoff dist: 400.0 and for *Satb1* cKO: upfreq 0.0, lowfreq –2.0, maxdist: 1100, cutoff dist: 800.0. We used the optimized set of parameters and based on correlation between the model and the experimental contact maps, we selected the top 5,000 models (out of 25,000 models) to generate a production ensemble. Production ensembles were well correlated with the input Hi-C contact matrices (Spearman’s r = 0.716 for WT and r = 0.7877 for *Satb1* cKO).

The production ensemble models were clustered using TADbit’s built-in objective function and the MCL Markov clustering (Enright et al., 2002). The centroid model of the first cluster for each genotype was visualized using the UCSF Chimera molecular visualization software (Pettersen et al., 2004) at a scale of 1:4. The models were colored according to the distance from the 5’-end. The highlighted beads represent an approximation of SATB1 loop anchors derived from SATB1 HiChIP data at *Bcl6* (yellow) and its super-enhancers (black). The WT models were rotated to obtain a clear view of all relevant structures and then aligned to the *Satb1* cKO-derived models using the match command.

The three-dimensional structures in the production ensemble were used to calculate the distance distribution between the beads corresponding to the loci closest to the *Bcl6* locus and its proximal super-enhancer 1 (corresponding to chr16:23980000-24000000 and chr16:24240000-24260000, respectively). The resulting distributions of the WT and *Satb1* cKO data were compared using the Mann-Whitney U test and assessed for multimodality with the skinny-dip test (Maurus and Plant, 2016). The distances between *Bcl6* and its super-enhancer based on the ensemble of the 5,000 sampled models were: mean rank WT: 2616.3162; mean rank *Satb1* cKO: 7384.6838 (p = 0.0; Mann-Whitney U test). Dip test results (distribution modes): WT: [171.205, 200.087], [200.248, 298.606] (p < 0.001); *Satb1* cKO: [198.063, 441.868].

## References

Alvarez, J.D., Yasui, D.H., Niida, H., Joh, T., Loh, D.Y., and Kohwi-Shigematsu, T. (2000). The MAR-binding protein SATB1 orchestrates temporal and spatial expression of multiple genes during T-cell development. Genes Dev. 14, 521–535.

Apostolou, E., Ferrari, F., Walsh, R.M., Bar-Nur, O., Stadtfeld, M., Cheloufi, S., Stuart, H.T., Polo, J.M., Ohsumi, T.K., Borowsky, M.L., et al. (2013). Genome-wide chromatin interactions of the Nanog locus in pluripotency, differentiation, and reprogramming. Cell Stem Cell 12, 699–712.

Bailey, T.L., and Elkan, C. (1994). Fitting a mixture model by expectation maximization to discover motifs in biopolymers. Proc. Int. Conf. Intell. Syst. Mol. Biol. 2, 28–36.

Balamotis, M.A., Tamberg, N., Woo, Y.J., Li, J., Davy, B., Kohwi-Shigematsu, T., and Kohwi, Y. (2012). Satb1 ablation alters temporal expression of immediate early genes and reduces dendritic spine density during postnatal brain development. Mol. Cell. Biol. 32, 333–347.

Bhattacharyya, S., Chandra, V., Vijayanand, P., and Ay, F. (2019). Identification of significant chromatin contacts from HiChIP data by FitHiChIP. Nat. Commun. 10, 1–14.

Bracken, A.P., and Helin, K. (2009). Polycomb group proteins: navigators of lineage pathways led astray in cancer. Nat. Rev. Cancer 9, 773–784.

Buus, T.B., Schmidt, J.D., Bonefeld, C.M., Geisler, C., and Lauritsen, J.P.H. (2016). Development of interleukin-17-producing Vγ2 + γδ T cells is reduced by ICOS signaling in the thymus. Oncotarget 7, 19341–19354.

Cai, S., Han, H.-J., and Kohwi-Shigematsu, T. (2003). Tissue-specific nuclear architecture and gene expession regulated by SATB1. Nat. Genet. 34, 42–51.

Cai, S., Lee, C.C., and Kohwi-Shigematsu, T. (2006). SATB1 packages densely looped, transcriptionally active chromatin for coordinated expression of cytokine genes. Nat. Genet. 38, 1278–1288.

Chapuy, B., McKeown, M.R., Lin, C.Y., Monti, S., Roemer, M.G.M., Qi, J., Rahl, P.B., Sun, H.H., Yeda, K.T., Doench, J.G., et al. (2013). Discovery and characterization of super-enhancer associated dependencies in diffuse large B-cell lymphoma. Cancer Cell 24, 777–790.

Chen, L., Carico, Z., Shih, H.-Y., and Krangel, M.S. (2015). A discrete chromatin loop in the mouse *Tcra* - *Tcrd* locus shapes the TCRδ and TCRα repertoires. Nat. Immunol. 16, 1085–1093.

Cho, W.-K., Spille, J.-H., Hecht, M., Lee, C., Li, C., Grube, V., and Cisse, I.I. (2018). Mediator and RNA polymerase II clusters associate in transcription-dependent condensates. Science 361, 412–415.

Corces, M.R., Trevino, A.E., Hamilton, E.G., Greenside, P.G., Sinnott-Armstrong, N.A., Vesuna, S., Satpathy, A.T., Rubin, A.J., Montine, K.S., Wu, B., et al. (2017). An improved ATAC-seq protocol reduces background and enables interrogation of frozen tissues. Nat. Methods 14, 959–962.

Cresswell, K.G., and Dozmorov, M.G. (2020). TADCompare: An R package for differential and temporal analysis of topologically associated domains. Front. Genet. 11.

Crump, N.T., Ballabio, E., Godfrey, L., Thorne, R., Repapi, E., Kerry, J., Tapia, M., Hua, P., Lagerholm, C., Filippakopoulos, P., et al. (2021). BET inhibition disrupts transcription but retains enhancer-promoter contact. Nat. Commun. 12, 223.

Davis, C.A., Hitz, B.C., Sloan, C.A., Chan, E.T., Davidson, J.M., Gabdank, I., Hilton, J.A., Jain, K., Baymuradov, U.K., Narayanan, A.K., et al. (2018). The Encyclopedia of DNA elements (ENCODE): data portal update. Nucleic Acids Res. 46, D794–D801.

Denaxa, M., Kalaitzidou, M., Garefalaki, A., Achimastou, A., Lasrado, R., Maes, T., and Pachnis, V. (2012). Maturation-promoting activity of SATB1 in MGE-derived cortical interneurons. Cell Rep. 2, 1351–1362.

Deng, W., Lee, J., Wang, H., Miller, J., Reik, A., Gregory, P.D., Dean, A., and Blobel, G.A. (2012). Controlling long-range genomic interactions at a native locus by targeted tethering of a looping factor. Cell 149, 1233–1244.

Dickinson, L.A., Joh, T., Kohwi, Y., and Kohwi-Shigematsu, T. (1992). A tissue-specific MAR/SAR DNA-binding protein with unusual binding site recognition. Cell 70, 631–645.

Dixon, J.R., Selvaraj, S., Yue, F., Kim, A., Li, Y., Shen, Y., Hu, M., Liu, J.S., and Ren, B. (2012). Topological domains in mammalian genomes identified by analysis of chromatin interactions. Nature 485, 376–380.

Dunham, I., Kundaje, A., Aldred, S.F., Collins, P.J., Davis, C.A., Doyle, F., Epstein, C.B., Frietze, S., Harrow, J., Kaul, R., et al. (2012). An integrated encyclopedia of DNA elements in the human genome. Nature 489, 57–74.

Durand, N.C., Shamim, M.S., Machol, I., Rao, S.S.P., Huntley, M.H., Lander, E.S., and Aiden, E.L. (2016). Juicer provides a one-click system for analyzing loop-resolution Hi-C experiments. Cell Syst. 3, 95–98.

Emmanuel, A.O., Arnovitz, S., Haghi, L., Mathur, P.S., Mondal, S., Quandt, J., Okoreeh, M.K., Maienschein-Cline, M., Khazaie, K., Dose, M., et al. (2018). TCF-1 and HEB cooperate to establish the epigenetic and transcription profiles of CD4^+^ CD8^+^ thymocytes. Nat. Immunol. 19, 1366–1378.

Enright, A.J., Van Dongen, S., and Ouzounis, C.A. (2002). An efficient algorithm for large-scale detection of protein families. Nucleic Acids Res. 30, 1575–1584.

Feng, D., Li, Z., Qin, L., and Hao, B. (2021). The role of chromatin organizer Satb1 in shaping TCR repertoire in adult thymus. Genome.

Fessing, M.Y., Mardaryev, A.N., Gdula, M.R., Sharov, A.A., Sharova, T.Y., Rapisarda, V., Gordon, K.B., Smorodchenko, A.D., Poterlowicz, K., Ferone, G., et al. (2011). p63 regulates *Satb1* to control tissue-specific chromatin remodeling during development of the epidermis. J. Cell Biol. 194, 825–839.

Fugmann, S.D., Lee, A.I., Shockett, P.E., Villey, I.J., and Schatz, D.G. (2000). The RAG proteins and V(D)J recombination: complexes, ends, and transposition. Annu. Rev. Immunol. 18, 495–527.

Fujii, Y., Kumatori, A., and Nakamura, M. (2003). SATB1 makes a complex with p300 and represses gp91phox promoter activity. Microbiol. Immunol. 47, 803–811.

Garcia-Perez, L., Famili, F., Cordes, M., Brugman, M., van Eggermond, M., Wu, H., Chouaref, J., Granado, D.S.L., Tiemessen, M.M., Pike-Overzet, K., et al. (2020). Functional definition of a transcription factor hierarchy regulating T cell lineage commitment. Sci. Adv. 6.

Ghosh, R.P., Shi, Q., Yallg, L., Reddick, M.P., Nikitina, T., Zhurkin, V.B., Fordyce, P., Stasevich, T.J., Chang, H.Y., Greenleaf, W.J., et al. (2019). Satb1 integrates DNA binding site geometry and torsional stress to differentially target nucleosome-dense regions. Nat. Commun. 10, 3221.

Giammartino, D.C.D., Polyzos, A., and Apostolou, E. (2020). Transcription factors: building hubs in the 3D space. Cell Cycle 19, 2395–2410.

Gioulbasani, M., Galaras, A., Grammenoudi, S., Moulos, P., Dent, A.L., Sigvardsson, M., Hatzis, P., Kee, B.L., and Verykokakis, M. (2020). The transcription factor BCL-6 controls early development of innate-like T cells. Nat. Immunol. 1–12.

Haas, J.D., Ravens, S., Düber, S., Sandrock, I., Oberdörfer, L., Kashani, E., Chennupati, V., Föhse, L., Naumann, R., Weiss, S., et al. (2012). Development of interleukin-17-producing γδ T cells is restricted to a functional embryonic wave. Immunity 37, 48–59.

Hao, B., Naik, A.K., Watanabe, A., Tanaka, H., Chen, L., Richards, H.W., Kondo, M., Taniuchi, I., Kohwi, Y., Kohwi-Shigematsu, T., et al. (2015). An anti-silencer- and SATB1-dependent chromatin hub regulates *Rag1* and *Rag2* gene expression during thymocyte development. J. Exp. Med. 212, 809–824.

Heinz, S., Benner, C., Spann, N., Bertolino, E., Lin, Y.C., Laslo, P., Cheng, J.X., Murre, C., Singh, H., and Glass, C.K. (2010). Simple combinations of lineage-determining transcription factors prime cis-regulatory elements required for macrophage and B cell identities. Mol. Cell 38, 576–589.

Hsieh, T.-H.S., Cattoglio, C., Slobodyanyuk, E., Hansen, A.S., Rando, O.J., Tjian, R., and Darzacq, X. (2020). Resolving the 3D landscape of transcription-linked mammalian chromatin folding. Mol. Cell 78, 539–553.e8.

Hu, G., Cuil, K., Fang, D., Hirose, S., Wang, X., Wangsa, D., Jin, W., Ried, T., Liu, P., Zhu, J., et al. (2018). Transformation of accessible chromatin and 3D nucleome underlies lineage commitment of early T cells. Immunity 48, 227–242.e8.

Hua, P., Badat, M., Hanssen, L.L.P., Hentges, L.D., Crump, N., Downes, D.J., Jeziorska, D.M., Oudelaar, A.M., Schwessinger, R., Taylor, S., et al. (2021). Defining genome architecture at base-pair resolution. Nature.

Hyjek, E., Chadburn, A., Liu, Y.F., Cesarman, E., and Knowles, D.M. (2001). BCL-6 protein is expressed in precursor T-cell lymphoblastic lymphoma and in prenatal and postnatal thymus. Blood 97, 270–276.

Ing-Simmons, E., Seitan, V., Faure, A., Flicek, P., Carroll, T., Dekker, J., Fisher, A., Lenhard, B., and Merkenschlager, M. (2015). Spatial enhancer clustering and regulation of enhancer-proximal genes by cohesin. Genome Res. gr.184986.114.

Jangid, R., Jayani, R., and Galande, S. (2014). Chromatin organizer SATB1 recruits Set9 histone methyltransferase to regulate global gene expression. Genes Genet. Syst. 89, 275–275.

Ji, Y., Resch, W., Corbett, E., Yamane, A., Casellas, R., and Schatz, D.G. (2010). The in vivo pattern of binding of RAG1 and RAG2 to antigen receptor loci. Cell 141, 419–431.

Johnson, J.L., Georgakilas, G., Petrovic, J., Kurachi, M., Cai, S., Harly, C., Pear, W.S., Bhandoola, A., Wherry, E.J., and Vahedi, G. (2018). Lineage-determining transcription factor TCF-1 initiates the epigenetic identity of T cell development. Immunity 48, 243–257.e10.

Johnston, R.J., Poholek, A.C., DiToro, D., Yusuf, I., Eto, D., Barnett, B., Dent, A.L., Craft, J., and Crotty, S. (2009). Bcl6 and Blimp-1 are reciprocal and antagonistic regulators of T follicular helper cell differentiation. Science 325, 1006–1010.

Kakugawa, K., Kojo, S., Tanaka, H., Seo, W., Endo, T.A., Kitagawa, Y., Muroi, S., Tenno, M., Yasmin, N., Kohwi, Y., et al. (2017). Essential roles of SATB1 in specifying T lymphocyte subsets. Cell Rep. 19, 1176–1188.

Kim, S., and Shendure, J. (2019). Mechanisms of interplay between transcription factors and the 3D genome. Mol. Cell 76, 306–319.

Kim, D., Paggi, J.M., Park, C., Bennett, C., and Salzberg, S.L. (2019). Graph-based genome alignment and genotyping with HISAT2 and HISAT-genotype. Nat. Biotechnol. 37, 907–915.

Kitagawa, Y., Ohkura, N., Kidani, Y., Vandenbon, A., Hirota, K., Kawakami, R., Yasuda, K., Motooka, D., Nakamura, S., Kondo, M., et al. (2017). Guidance of regulatory T cell development by Satb1-dependent super-enhancer establishment. Nat. Immunol. 18, 173–183.

Knight, P.A., and Ruiz, D. (2013). A fast algorithm for matrix balancing. IMA J. Numer. Anal. 33, 1029–1047.

Kohwi Shigematsu, T., Maass, K., and Bode, J. (1997). A thymocyte factor SATB1 suppresses transcription of stably integrated matrix-attachment region-linked reporter genes. Biochemistry 36, 12005–12010.

Kondo, M., Tanaka, Y., Kuwabara, T., Naito, T., Kohwi-Shigematsu, T., and Watanabe, A. (2016). SATB1 plays a critical role in establishment of immune tolerance. J. Immunol. 196, 563–572.

Kumar, P.P., Purbey, P.K., Ravi, D.S., Mitra, D., and Galande, S. (2005). Displacement of SATB1-bound histone deacetylase 1 corepressor by the human immunodeficiency virus type 1 transactivator induces expression of interleukin-2 and its receptor in T cells. Mol. Cell. Biol. 25, 1620–1633.

Kumar, P.P., Purbey, P.K., Sinha, C.K., Notani, D., Limaye, A., Jayani, R.S., and Galande, S. (2006). Phosphorylation of SATB1, a global gene regulator, acts as a molecular switch regulating its transcriptional activity in vivo. Mol. Cell 22, 231–243.

Langmead, B., and Salzberg, S.L. (2012). Fast gapped-read alignment with Bowtie 2. Nat. Methods 9, 357–359.

Li, H., Handsaker, B., Wysoker, A., Fennell, T., Ruan, J., Homer, N., Marth, G., Abecasis, G., Durbin, R., and Subgroup, 1000 Genome Project Data Processing (2009). The Sequence Alignment/Map format and SAMtools. Bioinformatics 25, 2078–2079.

Liao, Y., Smyth, G.K., and Shi, W. (2014). featureCounts: an efficient general purpose program for assigning sequence reads to genomic features. Bioinformatics 30, 923–930.

Lieberman-Aiden, E., Berkum, N.L. van, Williams, L., Imakaev, M., Ragoczy, T., Telling, A., Amit, I., Lajoie, B.R., Sabo, P.J., Dorschner, M.O., et al. (2009). Comprehensive mapping of long-range interactions reveals folding principles of the human genome. Science 326, 289–293.

Liu, J., Bramblett, D., Zhu, Q., Lozano, M., Kobayashi, R., Ross, S.R., and Dudley, J.P. (1997). The matrix attachment region-binding protein SATB1 participates in negative regulation of tissue-specific gene expression. Mol. Cell. Biol. 17, 5275–5287.

Lopes, N., Sergé, A., Ferrier, P., and Irla, M. (2015). Thymic crosstalk coordinates medulla organization and T-cell tolerance induction. Front. Immunol. 6.

Love, M.I., Huber, W., and Anders, S. (2014). Moderated estimation of fold change and dispersion for RNA-seq data with DESeq2. Genome Biol. 15, 550.

Lun, A.T.L., and Smyth, G.K. (2015). diffHic: a Bioconductor package to detect differential genomic interactions in Hi-C data. BMC Bioinformatics 16, 258.

Lupiáñez, D.G., Kraft, K., Heinrich, V., Krawitz, P., Brancati, F., Klopocki, E., Horn, D., Kayserili, H., Opitz, J.M., Laxova, R., et al. (2015). Disruptions of topological chromatin domains cause pathogenic rewiring of gene-enhancer interactions. Cell 161, 1012–1025.

Maman, Y., Teng, G., Seth, R., Kleinstein, S.H., and Schatz, D.G. (2016). RAG1 targeting in the genome is dominated by chromatin interactions mediated by the non-core regions of RAG1 and RAG2. Nucleic Acids Res. 44, 9624–9637.

Mathew, R., Mao, A., Chiang, A.H., Bertozzi-Villa, C., Bunker, J.J., Scanlon, S.T., McDonald, B.D., Constantinides, M.G., Hollister, K., Singer, J.D., et al. (2014). A negative feedback loop mediated by the Bcl6–cullin 3 complex limits Tfh cell differentiation. J. Exp. Med. 211, 1137–1151.

Maurus, S., and Plant, C. (2016). Skinny-dip: clustering in a sea of noise. In Proceedings of the 22nd ACM SIGKDD International Conference on Knowledge Discovery and Data Mining, (San Francisco California USA: ACM), pp. 1055–1064.

Mumbach, M.R., Rubin, A.J., Flynn, R.A., Dai, C., Khavari, P.A., Greenleaf, W.J., and Chang, H.Y. (2016). HiChIP: efficient and sensitive analysis of protein-directed genome architecture. Nat. Methods 13, 919–922.

Muñoz-Ruiz, M., Sumaria, N., Pennington, D.J., and Silva-Santos, B. (2017). Thymic determinants of γδ T cell differentiation. Trends Immunol. 38, 336–344.

Nora, E.P., Lajoie, B.R., Schulz, E.G., Giorgetti, L., Okamoto, I., Servant, N., Piolot, T., van Berkum, N.L., Meisig, J., Sedat, J., et al. (2012). Spatial partitioning of the regulatory landscape of the X-inactivation center. Nature 485, 381–385.

Nora, E.P., Goloborodko, A., Valton, A.-L., Gibcus, J.H., Uebersohn, A., Abdennur, N., Dekker, J., Mirny, L.A., and Bruneau, B.G. (2017). Targeted degradation of CTCF decouples local insulation of chromosome domains from genomic compartmentalization. Cell 169, 930–944.e22.

Nurieva, R.I., Chung, Y., Martinez, G.J., Yang, X.O., Tanaka, S., Matskevitch, T.D., Wang, Y.-H., and Dong, C. (2009). Bcl6 mediates the development of T follicular helper cells. Science 325, 1001–1005.

Oki, S., Ohta, T., Shioi, G., Hatanaka, H., Ogasawara, O., Okuda, Y., Kawaji, H., Nakaki, R., Sese, J., and Meno, C. (2018). ChIP-Atlas: a data-mining suite powered by full integration of public ChIP-seq data. EMBO Rep. 19, e46255.

Papotto, P.H., Ribot, J.C., and Silva-Santos, B. (2017). IL-17+ γδ T cells as kick-starters of inflammation. Nat. Immunol. 18, 604–611.

Pertea, M., Pertea, G.M., Antonescu, C.M., Chang, T.-C., Mendell, J.T., and Salzberg, S.L. (2015). StringTie enables improved reconstruction of a transcriptome from RNA-seq reads. Nat. Biotechnol. 33, 290–295.

Peters, J.-M. (2021). How DNA loop extrusion mediated by cohesin enables V(D)J recombination. Curr. Opin. Cell Biol. 70, 75–83.

Pettersen, E.F., Goddard, T.D., Huang, C.C., Couch, G.S., Greenblatt, D.M., Meng, E.C., and Ferrin, T.E. (2004). UCSF Chimera--a visualization system for exploratory research and analysis. J. Comput. Chem. 25, 1605–1612.

Phillips, J.E., and Corces, V.G. (2009). CTCF: Master weaver of the genome. Cell 137, 1194–1211.

Purbey, P.K., Singh, S., Notani, D., Kumar, P.P., Limaye, A.S., and Galande, S. (2009). Acetylation-dependent interaction of SATB1 and CtBP1 mediates transcriptional repression by SATB1. Mol. Cell. Biol. 29, 1321–1337.

Qian, J., Wang, Q., Dose, M., Pruett, N., Kieffer-Kwon, K.-R., Resch, W., Liang, G., Tang, Z., Mathé, E., Benner, C., et al. (2014). B cell super-enhancers and regulatory clusters recruit AID tumorigenic activity. Cell 159, 1524–1537.

Quinlan, A.R., and Hall, I.M. (2010). BEDTools: a flexible suite of utilities for comparing genomic features. Bioinformatics 26, 841–842.

Ramachandrareddy, H., Bouska, A., Shen, Y., Ji, M., Rizzino, A., Chan, W.C., and McKeithan, T.W. (2010). BCL6 promoter interacts with far upstream sequences with greatly enhanced activating histone modifications in germinal center B cells. Proc. Natl. Acad. Sci. 107, 11930–11935.

Ramírez, F., Ryan, D.P., Grüning, B., Bhardwaj, V., Kilpert, F., Richter, A.S., Heyne, S., Dündar, F., and Manke, T. (2016). deepTools2: a next generation web server for deep-sequencing data analysis. Nucleic Acids Res. 44, W160–165.

Ramírez, F., Bhardwaj, V., Arrigoni, L., Lam, K.C., Grüning, B.A., Villaveces, J., Habermann, B., Akhtar, A., and Manke, T. (2018). High-resolution TADs reveal DNA sequences underlying genome organization in flies. Nat. Commun. 9, 189.

Rao, S.S.P., Huntley, M.H., Durand, N.C., Stamenova, E.K., Bochkov, I.D., Robinson, J.T., Sanborn, A.L., Machol, I., Omer, A.D., Lander, E.S., et al. (2014). A 3D map of the human genome at kilobase resolution reveals principles of chromatin looping. Cell 159, 1665–1680.

Rao, S.S.P., Huang, S.-C., Glenn St Hilaire, B., Engreitz, J.M., Perez, E.M., Kieffer-Kwon, K.-R., Sanborn, A.L., Johnstone, S.E., Bascom, G.D., Bochkov, I.D., et al. (2017). Cohesin loss eliminates all loop domains. Cell 171, 305–320.e24.

Reimand, J., Kull, M., Peterson, H., Hansen, J., and Vilo, J. (2007). g:Profiler–a web-based toolset for functional profiling of gene lists from large-scale experiments. Nucleic Acids Res. 35, W193–200.

Robinson, M.D., McCarthy, D.J., and Smyth, G.K. (2010). edgeR: a Bioconductor package for differential expression analysis of digital gene expression data. Bioinformatics 26, 139–140.

Rogers, C.H., Mielczarek, O., and Corcoran, A.E. (2021). Dynamic 3D locus organization and its drivers underpin immunoglobulin recombination. Front. Immunol. 11.

Rowley, M.J., and Corces, V.G. (2018). Organizational principles of 3D genome architecture. Nat. Rev. Genet. 19, 789–800.

Rowley, M.J., Nichols, M.H., Lyu, X., Ando-Kuri, M., Rivera, I.S.M., Hermetz, K., Wang, P., Ruan, Y., and Corces, V.G. (2017). Evolutionarily conserved principles predict 3D chromatin organization. Mol. Cell 67, 837–852.e7.

Rowley, M.J., Poulet, A., Nichols, M., Bixler, B., Sanborn, A., Brouhard, E., Hermetz, K., Linsenbaum, H., Csankovszki, G., Aiden, E.L., et al. (2020). Analysis of Hi-C data using SIP effectively identifies loops in organisms from <I>C. elegans<\i> to mammals. Genome Res. gr.257832.119.

Ryan, R.J.H., Drier, Y., Whitton, H., Cotton, M.J., Kaur, J., Issner, R., Gillespie, S., Epstein, C.B., Nardi, V., Sohani, A.R., et al. (2015). Detection of enhancer-associated rearrangements reveals mechanisms of oncogene dysregulation in B-cell lymphoma. Cancer Discov. 5, 1058–1071.

Sabari, B.R., Dall’Agnese, A., Boija, A., Klein, I.A., Coffey, E.L., Shrinivas, K., Abraham, B.J., Hannett, N.M., Zamudio, A.V., Manteiga, J.C., et al. (2018). Coactivator condensation at super-enhancers links phase separation and gene control. Science 361, eaar3958.

Sakaguchi, S., Yamaguchi, T., Nomura, T., and Ono, M. (2008). Regulatory T cells and immune tolerance. Cell 133, 775–787.

Sauerwald, N., Singhal, A., and Kingsford, C. (2020). Analysis of the structural variability of topologically associated domains as revealed by Hi-C. NAR Genomics Bioinforma. 2.

Schep, A.N., Buenrostro, J.D., Denny, S.K., Schwartz, K., Sherlock, G., and Greenleaf, W.J. (2015). Structured nucleosome fingerprints enable high-resolution mapping of chromatin architecture within regulatory regions. Genome Res. 25, 1757–1770.

Schindelin, J., Arganda-Carreras, I., Frise, E., Kaynig, V., Longair, M., Pietzsch, T., Preibisch, S., Rueden, C., Saalfeld, S., Schmid, B., et al. (2012). Fiji: an open-source platform for biological-image analysis. Nat. Methods 9, 676–682.

Schwarzer, W., Abdennur, N., Goloborodko, A., Pekowska, A., Fudenberg, G., Loe-Mie, Y., Fonseca, N.A., Huber, W., Haering, C., Mirny, L., et al. (2017). Two independent modes of chromatin organization revealed by cohesin removal. Nature 551, 51–56.

Seitan, V.C., Hao, B., Tachibana-Konwalski, K., Lavagnolli, T., Mira-Bontenbal, H., Brown, K.E., Teng, G., Carroll, T., Terry, A., Horan, K., et al. (2011). A role for cohesin in T cell receptor rearrangement and thymocyte differentiation. Nature 476, 467–471.

Seitan, V.C., Faure, A.J., Zhan, Y., McCord, R.P., Lajoie, B.R., Ing-Simmons, E., Lenhard, B., Giorgetti, L., Heard, E., Fisher, A.G., et al. (2013). Cohesin-based chromatin interactions enable regulated gene expression within preexisting architectural compartments. Genome Res. 23, 2066–2077.

Seo, J., Lozano, M.M., and Dudley, J.P. (2005). Nuclear matrix binding regulates SATB1-mediated transcriptional repression. J. Biol. Chem. 280, 24600–24609.

Serra, F., Baù, D., Goodstadt, M., Castillo, D., Filion, G.J., and Marti-Renom, M.A. (2017). Automatic analysis and 3D-modelling of Hi-C data using TADbit reveals structural features of the fly chromatin colors. PLOS Comput. Biol. 13, e1005665.

Servant, N., Varoquaux, N., Lajoie, B.R., Viara, E., Chen, C.-J., Vert, J.-P., Heard, E., Dekker, J., and Barillot, E. (2015). HiC-Pro: an optimized and flexible pipeline for Hi-C data processing. Genome Biol. 16, 259.

Shen, Y., Yue, F., McCleary, D.F., Ye, Z., Edsall, L., Kuan, S., Wagner, U., Dixon, J., Lee, L., Lobanenkov, V.V., et al. (2012). A map of the cis -regulatory sequences in the mouse genome. Nature 488, 116–120.

Shih, H.-Y., Verma-Gaur, J., Torkamani, A., Feeney, A.J., Galjart, N., and Krangel, M.S. (2012). *Tcra* gene recombination is supported by a *Tcra* enhancer- and CTCF-dependent chromatin hub. Proc. Natl. Acad. Sci. U. S. A. 109, E3493–3502.

Spilianakis, C.G., and Flavell, R.A. (2004). Long-range intrachromosomal interactions in the T helper type 2 cytokine locus. Nat. Immunol. 5, 1017–1027.

Spilianakis, C.G., Lalioti, M.D., Town, T., Lee, G.R., and Flavell, R.A. (2005). Interchromosomal associations between alternatively expressed loci. Nature 435, 637–645.

Stadhouders, R., Vidal, E., Serra, F., Di Stefano, B., Le Dily, F., Quilez, J., Gomez, A., Collombet, S., Berenguer, C., Cuartero, Y., et al. (2018). Transcription factors orchestrate dynamic interplay between genome topology and gene regulation during cell reprogramming. Nat. Genet. 50, 238–249.

Stadhouders, R., Filion, G.J., and Graf, T. (2019). Transcription factors and 3D genome conformation in cell-fate decisions. Nature 569, 345–354.

Sun, Z., Unutmaz, D., Zou, Y.-R., Sunshine, M.J., Pierani, A., Brenner-Morton, S., Mebius, R.E., and Littman, D.R. (2000). Requirement for RORγ in thymocyte survival and lymphoid organ development. Science 288, 2369–2373.

Sunkara, K.P., Gupta, G., Hansbro, P.M., Dua, K., and Bebawy, M. (2018). Functional relevance of SATB1 in immune regulation and tumorigenesis. Biomed. Pharmacother. 104, 87–93.

Teng, G., Maman, Y., Resch, W., Kim, M., Yamane, A., Qian, J., Kieffer-Kwon, K.-R., Mandal, M., Ji, Y., Meffre, E., et al. (2015). RAG represents a widespread threat to the lymphocyte genome. Cell 162, 751–765.

Vinuesa, C.G., Linterman, M.A., Yu, D., and MacLennan, I.C.M. (2016). Follicular helper T cells. Annu. Rev. Immunol. 34, 335–368.

Wang, Z., Yang, X., Chu, X., Zhang, J., Zhou, H., Shen, Y., and Long, J. (2012). The structural basis for the oligomerization of the N-terminal domain of SATB1. Nucleic Acids Res. 40, 4193–4202.

Wang, Z., Yang, X., Guo, S., Yang, Y., Su, X.-C., Shen, Y., and Long, J. (2014). Crystal structure of the ubiquitin-like domain-CUT repeat-like tandem of special AT-rich sequence binding protein 1 (SATB1) reveals a coordinating DNA-binding mechanism. J. Biol. Chem. 289, 27376–27385.

Wei, Z., Zhang, W., Fang, H., Li, Y., and Wang, X. (2018). esATAC: an easy-to-use systematic pipeline for ATAC-seq data analysis. Bioinformatics 34, 2664–2665.

Weintraub, A.S., Li, C.H., Zamudio, A.V., Sigova, A.A., Hannett, N.M., Day, D.S., Abraham, B.J., Cohen, M.A., Nabet, B., Buckley, D.L., et al. (2017). YY1 is a structural regulator of enhancer-promoter loops. Cell 171, 1573–1588.e28.

Wu, H., Deng, Y., Zhao, M., Zhang, J., Zheng, M., Chen, G., Li, L., He, Z., and Lu, Q. (2018). Molecular control of follicular helper T cell development and differentiation. Front. Immunol. 9.

Yasui, D., Miyano, M., Cai, S.T., Varga-WEisz, P., and Kohwi-Shigematsu, T. (2002). SATB1 targets chromatin remodelling to regulate genes over long distances. Nature 419, 641–645.

Yu, D., Rao, S., Tsai, L.M., Lee, S.K., He, Y., Sutcliffe, E.L., Srivastava, M., Linterman, M., Zheng, L., Simpson, N., et al. (2009). The transcriptional repressor Bcl-6 directs T follicular helper cell lineage commitment. Immunity 31, 457–468.

Zelenka, T., and Spilianakis, C. (2020). SATB1-mediated chromatin landscape in T cells. Nucleus 11, 117–131.

Zelenka, T., Tzerpos, P., Panagopoulos, G., Tsolis, K.C., Papamatheakis, D., Papadakis, V.M., Stanek, D., and Spilianakis, C.G. (Submitted). Physiological importance of SATB1 phase transitions and means of their regulation.

Zhao, H., Sun, Z., Wang, J., Huang, H., Kocher, J.-P., and Wang, L. (2014). CrossMap: a versatile tool for coordinate conversion between genome assemblies. Bioinforma. Oxf. Engl. 30, 1006–1007.

Zuberbuehler, M.K., Parker, M.E., Wheaton, J.D., Espinosa, J.R., Salzler, H.R., Park, E., and Ciofani, M. (2019). The transcription factor c-Maf is essential for the commitment of IL-17-producing γδ T cells. Nat. Immunol. 20, 73–85.

