## Supplemental Information for "The 3D enhancer network of the developing T cell genome is controlled by SATB1"

**Figure S1.**

(A) Phenotypical comparison of WT and *Satb1* cKO mice (SKO).

(B) Characterization of cell populations in WT and *Satb1* cKO by flow cytometry. Used animals were divided into young (45±11 days; 6 WT, 6 *Satb1* cKO) and old (179±35 days; 7 WT, 9 *Satb1* cKO) age categories. Only young animals were used for analysis of thymus due to its deterioration in old animals. Three young animals for each genotype were used for CD62 / CD44 analysis.

(C) *Satb1* cKO pancreas and lung sections were incubated with serum from either WT or *Satb1* cKO animals to detect the presence of autoantibodies. Scale bar in all images 100 μm.

(D) Differential analysis of topologically associating domains (TADs) between WT and *Satb1* cKO at 100 kbp resolution (combined biological replicates) showed high proportion of unchanged domains in *Satb1* cKO cells. As a control we provided differential analysis between two biological replicates, separately for WT and *Satb1* cKO samples. Black dots represent the distribution of different TAD categories for individual chromosomes.

(E) Aggregate peak analysis (APA; Rao et al., 2014) was calculated and visualized by Juicer Tools (Durand et al., 2016) to show that SATB1 HiChIP loops had stronger signal in WT Hi-C datasets than in *Satb1* cKO Hi-C. CTCF HiChIP loops retained the same APA score indicating that CTCF-based high order chromatin organization remained unchanged in *Satb1* cKO.

(F) Transcriptional insulation scores calculated as described in the methods section showed that expression of genes inside SATB1 and CTCF loops was different from genes outside the loops. Both SATB1 and CTCF evinced a similar insulation effect which was significantly different ($P < 2.22e{-16}$) between the loops and randomly shuffled loops in both WT and *Satb1* cKO. *P* values by Wilcoxon rank sum test.

(G) Analysis of A/B compartments performed according to the original protocol (Lieberman-Aiden et al., 2009) at 100,000 resolution of Hi-C datasets using HOMER (Heinz et al., 2010). Principle components of WT and *Satb1* cKO annotated regions were plotted against each other to show the proportion of compartment switch.

(H) Volcano plot of differentially accessible regions in WT and *Satb1* cKO as determined by ATAC-seq.

(I) Log2 fold change of ATAC-seq differential chromatin accessibility between WT and *Satb1* cKO plotted along genes ± 4 kbp. Graph depicts the drop in chromatin accessibility in *Satb1* cKO at the TSS of genes, supporting a regulatory role for SATB1.

(J) Correlation between RNA level and promoter chromatin accessibility changes (–2 kbp – TSS) of *Satb1* cKO differentially expressed genes (Spearman's $\rho = 0.438$, $P < 2.22e{-16}$, calculated based on correlation coefficients from 100 randomly selected sets of genes with mean Spearman's $\rho = 0.134$).

(K) Regions with low chromatin accessibility in WT had mostly increased chromatin accessibility in *Satb1* cKO. The average of log10 transformed read-normalized accessibility scores of ten randomizations depicted in Figure 3b was used as a cutoff to determine SATB1 peaks with low accessibility levels.

(L) Overlap enrichment between loop anchors and enhancers, presented separately for SATB1- and CTCF-mediated loops.

In (B and D), the horizontal lines inside violins represent the $25^{th}$, $50^{th}$ and $75^{th}$ percentiles. The red circle represents the mean ± s.d. *P* values by Wilcoxon rank sum test.

**Figure S2.**

(A) Diagnostic plot for residuals vs fitted values. Non-linear relationships between predictor variables and logFC changes were evaluated. Values were distributed randomly across a horizontal line with no obvious pattern arising, indicating that assumptions of the regression model were fulfilled well.

(B) Q-Q plot of residuals. The plot evaluates whether the residuals were normally distributed. Deviations from the normal distribution were observed for a set of residuals.

(C) Diagnostic plot for homoscedasticity. Model assumptions are not violated if data points are randomly distributed. No clear patterns were observed, indicating a good fit to the model assumptions.

(D) Diagnostic plot for influential outliers based on the Cook's distance. No gene was positioned outside the dashed line corresponding to the Cook's distance of 1. As such no gene was excluded when constructing the model.

(E) Evaluation of the quality of each predictor. A regression model was first constructed using all the predictors. For each predictor, a new model was constructed utilizing all predictors beside the one studied. The Akaike Information Criterion (AIC) of the old model was subtracted from the AIC of the new model. An increase in AIC after the removal of a predictor indicated that the model lacking that predictor performed worse than the original one. The final model included only the predictors that were associated with an increase in AIC after their removal.

(F) Coefficients of final predictors. Positive values indicate that a variable was linked with an increase in RNA levels in the *Satb1* cKO cells, while negative values indicate the opposite.

**Figure S3.**

(A) 3D computational modeling of the *Bcl6* locus utilizing chromatin contact domains (CCD) from HiChIP data. The model indicated that CTCF is responsible for maintaining the high order structure which is not sufficient to mediate the contact between the *Bcl6* gene and its enhancers. The combination of CTCF and SATB1-based models emphasized the importance of SATB1 in mediating the promoter-enhancer contacts. *Bcl6* enhancers are located in the green color-coded regions.

(B) Hi-C-derived 3D models for WT and *Satb1* cKO thymocytes with overlaid WT ChIP-seq data for histone modifications H3K27ac, H3K4me1 and H3K4me3 (visualized as a color-gradient). Models indicate how active enhancers decorated by H3K27ac and H3K4me1 are located in a close spatial proximity to *Bcl6* gene via SATB1-mediated chromatin interactions in WT and not in *Satb1* cKO cells. WT ChIP-seq data were used to emphasize the importance of the 3D organization. 1D H3K27ac ChIP-seq peaks derived from HiChIP experiments available for WT and *Satb1* cKO did not reveal any major differences between the genotypes, further reinforcing the importance of SATB1-mediated 3D chromatin organization regulating *Bcl6* expression. Position of beads corresponding to SATB1 loop anchors and demarcating the super-enhancer regions were: chr16:23985000-23990000 (*Bcl6*), chr16:24245000-24250000 (SE1) and chr16:24505000-24510000 (SE2).

(C) Quantified distances between Bcl6 and its super-enhancer 1 based on 5,000 Hi-C models in WT and in *Satb1* cKO. Mean rank WT: 2616.3162. Mean rank *Satb1* cKO: 7384.6838, $P = 0.0$ (Mann-Whitney U test), Dip test results: WT: [171.205, 200.087], [200.248, 298.606] and *Satb1* cKO: [198.063, 441.868].

**Figure S4.**

Genomic tracks and SATB1 HiChIP loops for the indicated important immune-related genes that evinced strong SATB1 looping connecting the genes with enhancers. The bar graph depicts the decrease in RNA expression values of selected genes in *Satb1* cKO thymocytes (log scale). Due to limited space, genes *Ets2* and *Socs2* which also demonstrate nice examples of SATB1-mediated regulatory loops were not included in the figure. Genes related to cellular adhesion and communication *Ccr7*, *Lta*, *Ltb* (including *Tnf*) are presented in Figure S6.

**Figure S5.**

(A) Gene ontology (GO) pathways of *Satb1* cKO underexpressed genes.

(B) GO pathways of *Satb1* cKO overexpressed genes.

(C) GO pathways of genes located in regions with less accessible chromatin in *Satb1* cKO.

(D) GO pathways of genes located in regions with more accessible chromatin in *Satb1* cKO.

(E) GO pathways of genes located in anchors of SATB1-mediated loops which were also H3K27ac underinteracting in *Satb1* cKO.

(F) GO pathways of genes located in anchors of SATB1-mediated loops which are also H3K27ac overinteracting in *Satb1* cKO.

**Figure S6.**

(A) Genomic tracks and SATB1 HiChIP loops indicating that the SATB1-mediated regulatory looping affect the *Tnf* locus.

(B) SATB1-mediated loops were also present at the *Ccr7* gene locus.

### References


Durand, N.C., Shamim, M.S., Machol, I., Rao, S.S.P., Huntley, M.H., Lander, E.S., and Aiden, E.L. (2016). Juicer provides a one-click system for analyzing loop-resolution Hi-C experiments. Cell Syst. *3*, 95–98.

Heinz, S., Benner, C., Spann, N., Bertolino, E., Lin, Y.C., Laslo, P., Cheng, J.X., Murre, C., Singh, H., and Glass, C.K. (2010). Simple combinations of lineage-determining transcription factors prime cis-regulatory elements required for macrophage and B cell identities. Mol. Cell *38*, 576–589.

Lieberman-Aiden, E., Berkum, N.L. van, Williams, L., Imakaev, M., Ragoczy, T., Telling, A., Amit, I., Lajoie, B.R., Sabo, P.J., Dorschner, M.O., et al. (2009). Comprehensive mapping of long-range interactions reveals folding principles of the human genome. Science *326*, 289–293.

Rao, S.S.P., Huntley, M.H., Durand, N.C., Stamenova, E.K., Bochkov, I.D., Robinson, J.T., Sanborn, A.L., Machol, I., Omer, A.D., Lander, E.S., et al. (2014). A 3D map of the human genome at kilobase resolution reveals principles of chromatin looping. Cell *159*, 1665–1680.

Figure S1

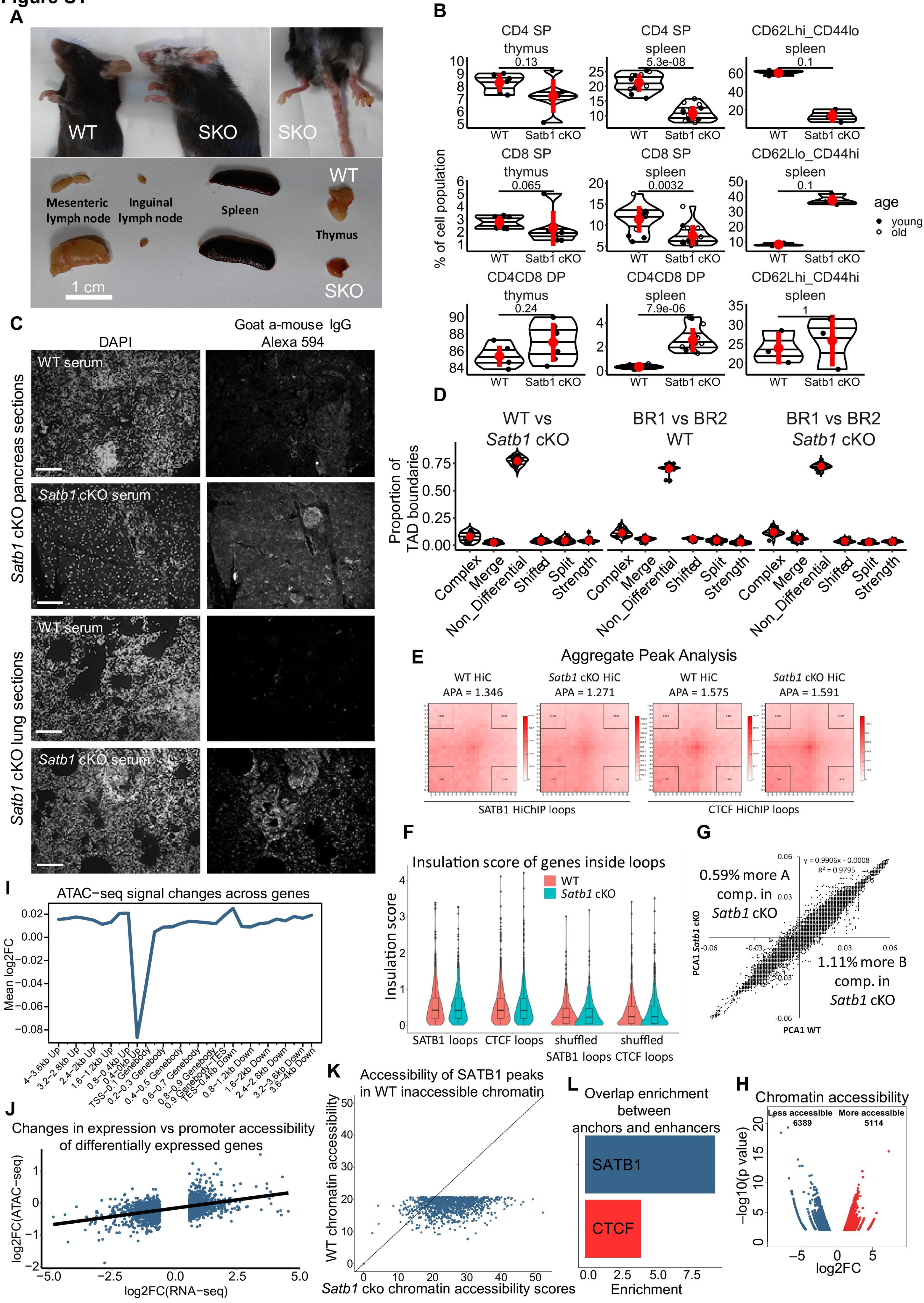
A
WT
SKO
SKO
WT
Mesenteric lymph node
Inguinal lymph node
Spleen
Thymus
1 cm
SKO
B
CD4 SP thymus
0.13
CD4 SP spleen
5.3e-08
CD62Lhi_CD44lo spleen
0.1
CD8 SP thymus
0.065
CD8 SP spleen
0.0032
CD62Llo_CD44hi spleen
0.1
CD4CD8 DP thymus
0.24
CD4CD8 DP spleen
7.9e-06
CD62Lhi_CD44hi spleen
1
% of cell population
WT
Satb1 cKO
age
young
old
C
DAPI
Goat a-mouse IgG Alexa 594
Satb1 cKO pancreas sections
WT serum
Satb1 cKO serum
Satb1 cKO lung sections
WT serum
Satb1 cKO serum
D
WT vs Satb1 cKO
BR1 vs BR2 WT
BR1 vs BR2 Satb1 cKO
Proportion of TAD boundaries
Complex
Merge
Non_Differential
Shifted
Split
Strength
E
Aggregate Peak Analysis
WT HiC APA = 1.346
Satb1 cKO HiC APA = 1.271
WT HiC APA = 1.575
Satb1 cKO HiC APA = 1.591
SATB1 HiChIP loops
CTCF HiChIP loops
F
Insulation score of genes inside loops
WT
Satb1 cKO
Insulation score
SATB1 loops
CTCF loops
shuffled SATB1 loops
shuffled CTCF loops
G
y = 0.9906x - 0.0008
R² = 0.9795
0.59% more A comp. in Satb1 cKO
1.11% more B comp. in Satb1 cKO
PCA1 Satb1 cKO
PCA1 WT
I
ATAC−seq signal changes across genes
Mean log2FC
4−3.6kb Up
3.2−2.8kb Up
2.4−2kb Up
1.6−1.2kb Up
0.8−0.4kb Up
0.4−0kb Up
TSS−0.1 Genebody
0.2−0.3 Genebody
0.4−0.5 Genebody
0.6−0.7 Genebody
0.8−0.9 Genebody
0.9 Genebody−TES
TES−0.4kb Down
0.8−1.2kb Down
1.6−2kb Down
2.4−2.8kb Down
3.2−3.6kb Down
3.6−4kb Down
J
Changes in expression vs promoter accessibility of differentially expressed genes
log2FC(ATAC−seq)
log2FC(RNA−seq)
K
Accessibility of SATB1 peaks in WT inaccessible chromatin
WT chromatin accessibility
Satb1 cko chromatin accessibility scores
L
Overlap enrichment between anchors and enhancers
SATB1
CTCF
Enrichment
H
Chromatin accessibility
Less accessible 6389
More accessible 5114
−log10(p value)
log2FC

**Figure S2**

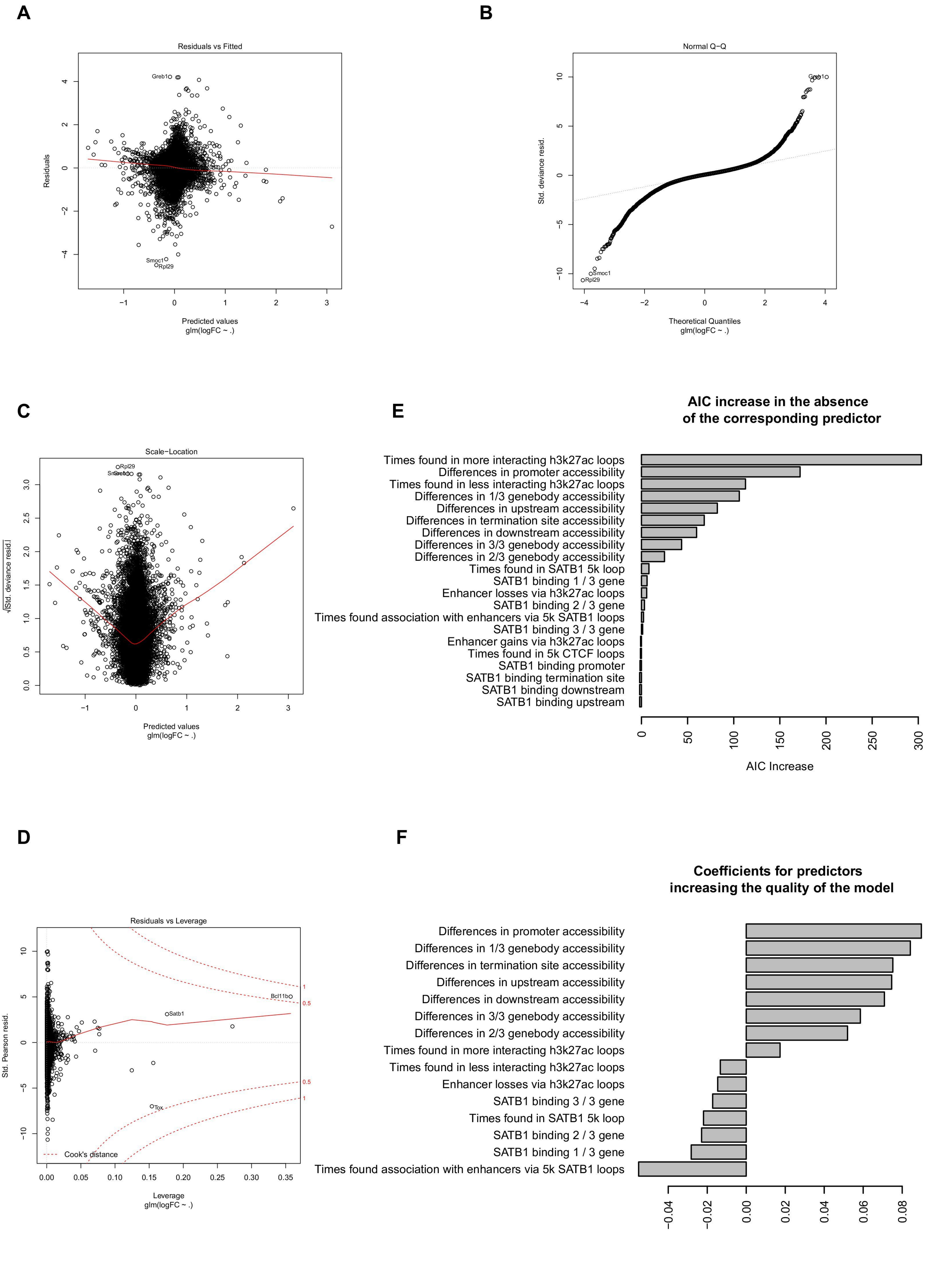
A
Residuals vs Fitted
Residuals
Predicted values
glm(logFC ~ .)
Greb1
Smoc1
Rpl29
B
Normal Q−Q
Std. deviance resid.
Theoretical Quantiles
glm(logFC ~ .)
Greb1
Smoc1
Rpl29
C
Scale−Location
√|Std. deviance resid.|
Predicted values
glm(logFC ~ .)
Rpl29
D
Residuals vs Leverage
Std. Pearson resid.
Leverage
glm(logFC ~ .)
Cook's distance
Bcl11b
Satb1
Tox
E
AIC increase in the absence of the corresponding predictor
Times found in more interacting h3k27ac loops
Differences in promoter accessibility
Times found in less interacting h3k27ac loops
Differences in 1/3 genebody accessibility
Differences in upstream accessibility
Differences in termination site accessibility
Differences in downstream accessibility
Differences in 3/3 genebody accessibility
Differences in 2/3 genebody accessibility
Times found in SATB1 5k loop
SATB1 binding 1 / 3 gene
Enhancer losses via h3k27ac loops
SATB1 binding 2 / 3 gene
Times found association with enhancers via 5k SATB1 loops
SATB1 binding 3 / 3 gene
Enhancer gains via h3k27ac loops
Times found in 5k CTCF loops
SATB1 binding promoter
SATB1 binding termination site
SATB1 binding downstream
SATB1 binding upstream
AIC Increase
F
Coefficients for predictors increasing the quality of the model
Differences in promoter accessibility
Differences in 1/3 genebody accessibility
Differences in termination site accessibility
Differences in upstream accessibility
Differences in downstream accessibility
Differences in 3/3 genebody accessibility
Differences in 2/3 genebody accessibility
Times found in more interacting h3k27ac loops
Times found in less interacting h3k27ac loops
Enhancer losses via h3k27ac loops
SATB1 binding 3 / 3 gene
Times found in SATB1 5k loop
SATB1 binding 2 / 3 gene
SATB1 binding 1 / 3 gene
Times found association with enhancers via 5k SATB1 loops

**Figure S3**

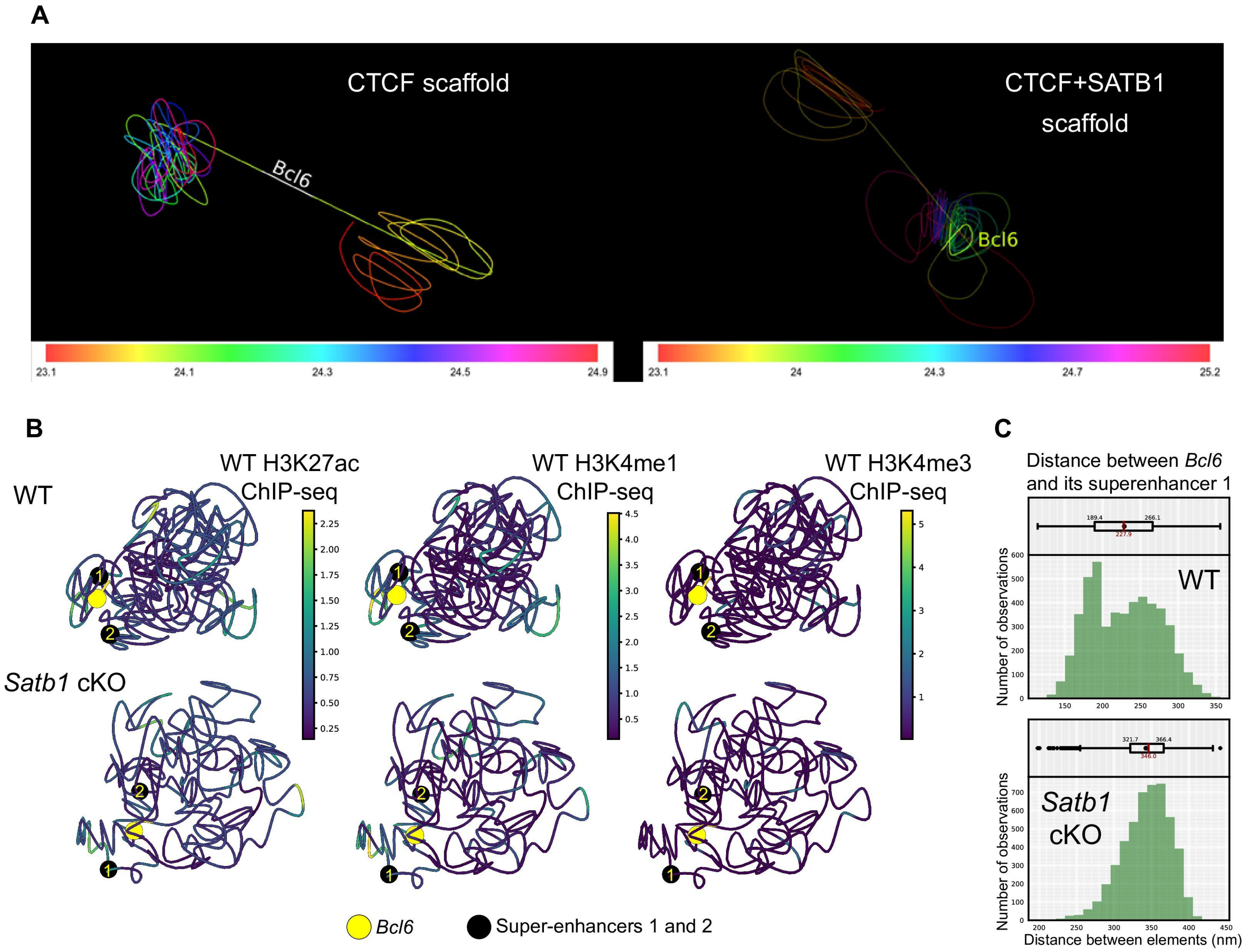
A
CTCF scaffold
Bcl6
CTCF+SATB1
scaffold
Bcl6
23.1
24.1
24.3
24.5
24.9
23.1
24
24.3
24.7
25.2
B
WT
WT H3K27ac
ChIP-seq
WT H3K4me1
ChIP-seq
WT H3K4me3
ChIP-seq
Satb1 cKO
Bcl6
Super-enhancers 1 and 2
C
Distance between Bcl6
and its superenhancer 1
WT
Satb1
cKO
Number of observations
Distance between elements (nm)

### Figure S4

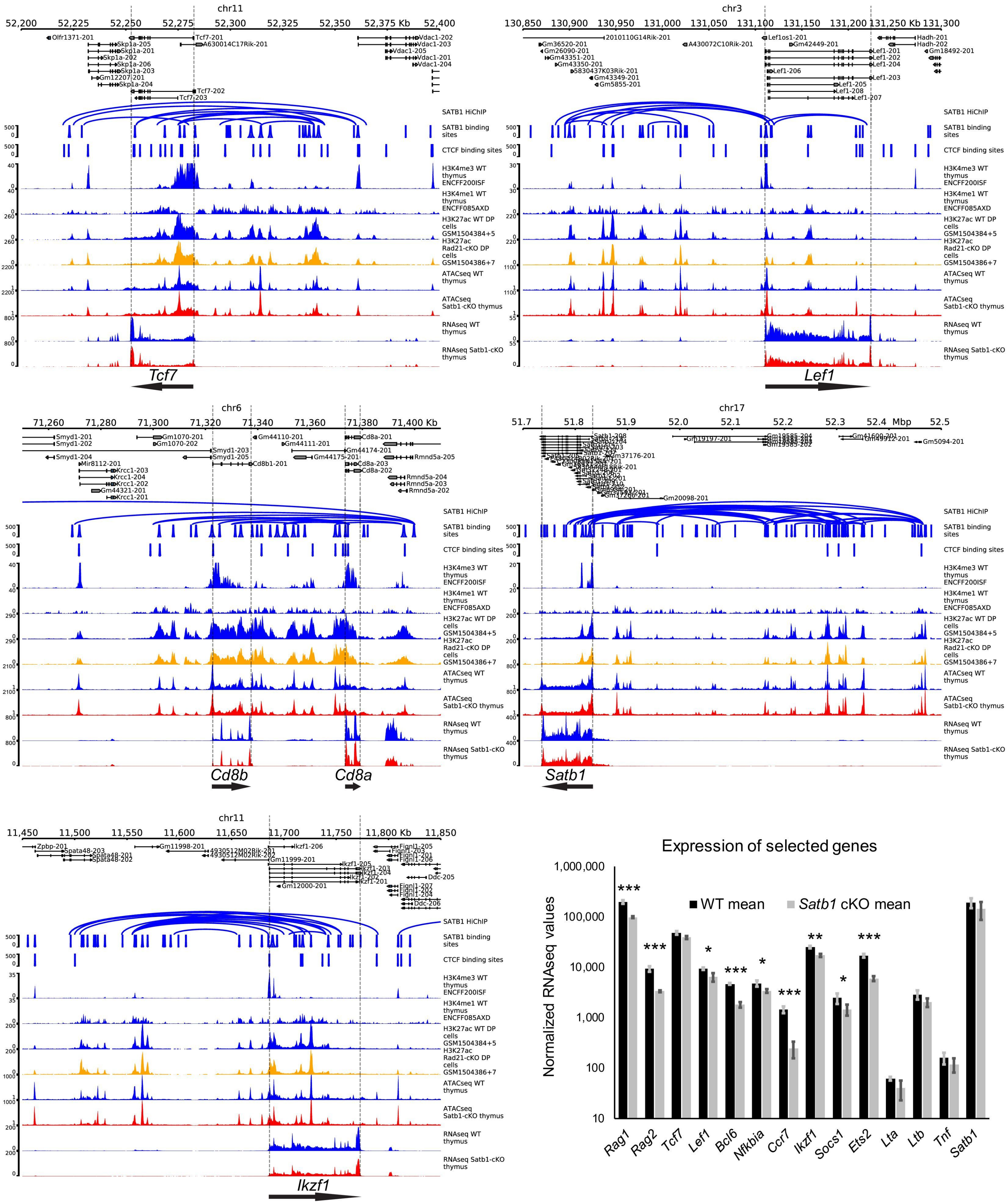

chr11
chr3
chr6
chr17
SATB1 HiChIP
SATB1 binding sites
CTCF binding sites
H3K4me3 WT thymus ENCFF200ISF
H3K4me1 WT thymus ENCFF085AXD
H3K27ac WT DP cells GSM1504384+5
H3K27ac Rad21-cKO DP cells GSM1504386+7
ATACseq WT thymus
ATACseq Satb1-cKO thymus
RNAseq WT thymus
RNAseq Satb1-cKO thymus
Tcf7
Lef1
Cd8b
Cd8a
Satb1
Ikzf1
Expression of selected genes
Normalized RNAseq values
WT mean
Satb1 cKO mean
Rag1
Rag2
Tcf7
Lef1
Bcl6
Nfkbia
Ccr7
Ikzf1
Socs1
Ets2
Lta
Ltb
Tnf
Satb1

**Figure S5**

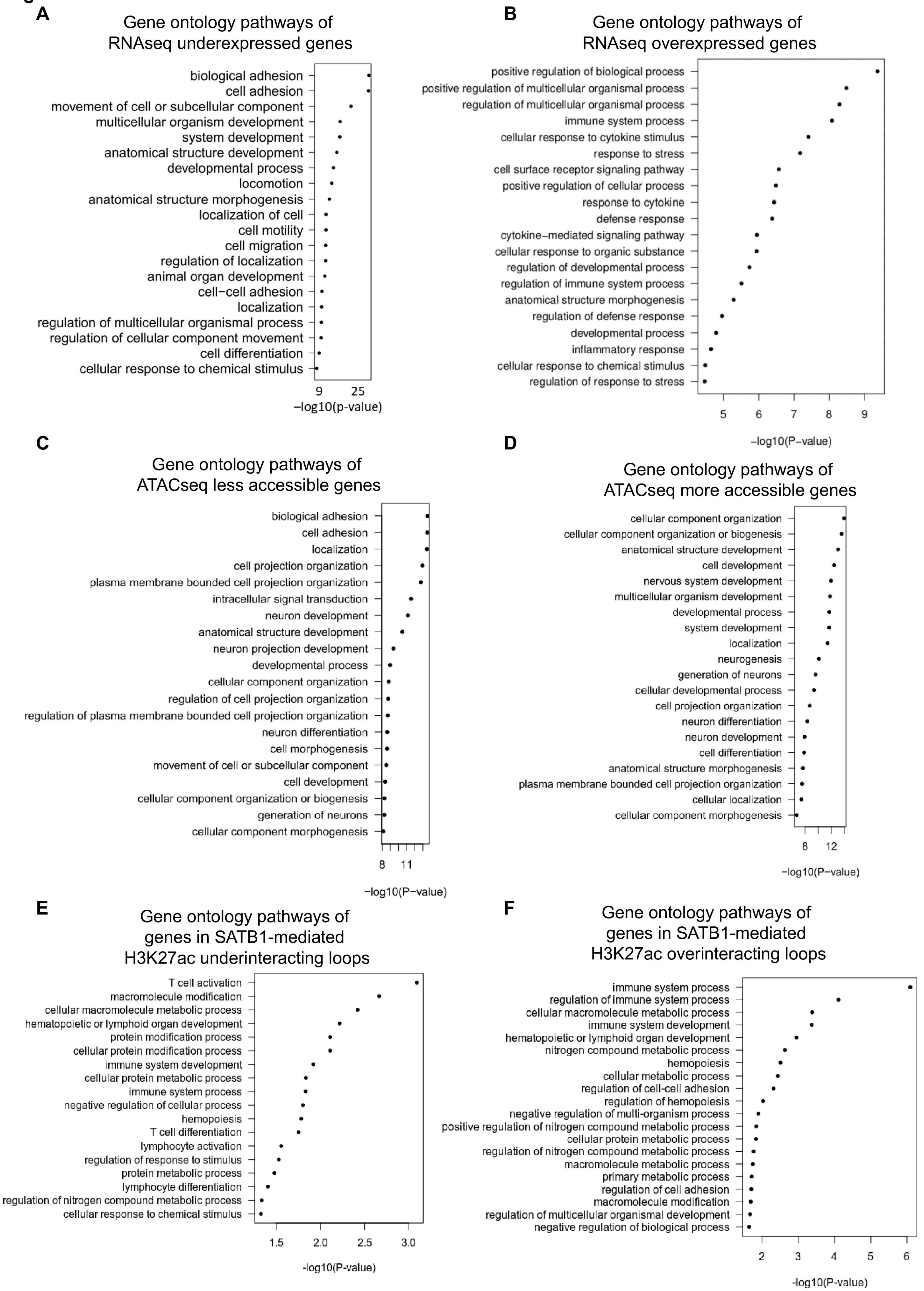

A
Gene ontology pathways of RNAseq underexpressed genes
biological adhesion
cell adhesion
movement of cell or subcellular component
multicellular organism development
system development
anatomical structure development
developmental process
locomotion
anatomical structure morphogenesis
localization of cell
cell motility
cell migration
regulation of localization
animal organ development
cell−cell adhesion
localization
regulation of multicellular organismal process
regulation of cellular component movement
cell differentiation
cellular response to chemical stimulus
9
25
−log10(p-value)
B
Gene ontology pathways of RNAseq overexpressed genes
positive regulation of biological process
positive regulation of multicellular organismal process
regulation of multicellular organismal process
immune system process
cellular response to cytokine stimulus
response to stress
cell surface receptor signaling pathway
positive regulation of cellular process
response to cytokine
defense response
cytokine−mediated signaling pathway
cellular response to organic substance
regulation of developmental process
regulation of immune system process
anatomical structure morphogenesis
regulation of defense response
developmental process
inflammatory response
cellular response to chemical stimulus
regulation of response to stress
5
6
7
8
9
−log10(P−value)
C
Gene ontology pathways of ATACseq less accessible genes
biological adhesion
cell adhesion
localization
cell projection organization
plasma membrane bounded cell projection organization
intracellular signal transduction
neuron development
anatomical structure development
neuron projection development
developmental process
cellular component organization
regulation of cell projection organization
regulation of plasma membrane bounded cell projection organization
neuron differentiation
cell morphogenesis
movement of cell or subcellular component
cell development
cellular component organization or biogenesis
generation of neurons
cellular component morphogenesis
8
11
−log10(P−value)
D
Gene ontology pathways of ATACseq more accessible genes
cellular component organization
cellular component organization or biogenesis
anatomical structure development
cell development
nervous system development
multicellular organism development
developmental process
system development
localization
neurogenesis
generation of neurons
cellular developmental process
cell projection organization
neuron differentiation
neuron development
cell differentiation
anatomical structure morphogenesis
plasma membrane bounded cell projection organization
cellular localization
cellular component morphogenesis
8
12
−log10(P−value)
E
Gene ontology pathways of genes in SATB1-mediated H3K27ac underinteracting loops
T cell activation
macromolecule modification
cellular macromolecule metabolic process
hematopoietic or lymphoid organ development
protein modification process
cellular protein modification process
immune system development
cellular protein metabolic process
immune system process
negative regulation of cellular process
hemopoiesis
T cell differentiation
lymphocyte activation
regulation of response to stimulus
protein metabolic process
lymphocyte differentiation
regulation of nitrogen compound metabolic process
cellular response to chemical stimulus
1.5
2.0
2.5
3.0
-log10(P-value)
F
Gene ontology pathways of genes in SATB1-mediated H3K27ac overinteracting loops
immune system process
regulation of immune system process
cellular macromolecule metabolic process
immune system development
hematopoietic or lymphoid organ development
nitrogen compound metabolic process
hemopoiesis
cellular metabolic process
regulation of cell-cell adhesion
regulation of hemopoiesis
negative regulation of multi-organism process
positive regulation of nitrogen compound metabolic process
cellular protein metabolic process
regulation of nitrogen compound metabolic process
macromolecule metabolic process
primary metabolic process
regulation of cell adhesion
macromolecule modification
regulation of multicellular organismal development
negative regulation of biological process
2
3
4
5
6
-log10(P-value)

**Figure S6**

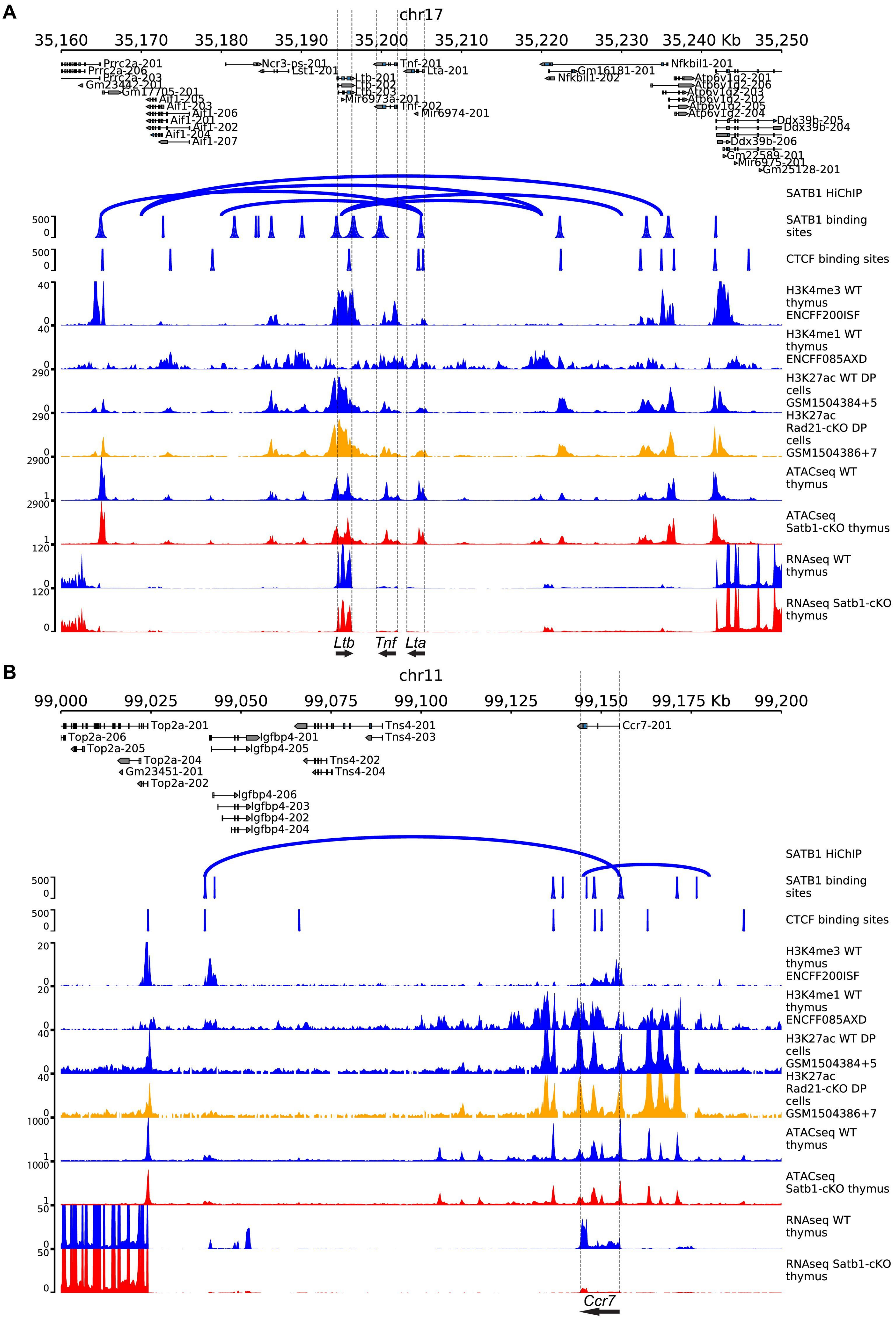
A
chr17
35,160
35,170
35,180
35,190
35,200
35,210
35,220
35,230
35,240 Kb
35,250
Prrc2a-201
Prrc2a-206
Prrc2a-203
Gm23442-201
Gm17705-201
Aif1-205
Aif1-203
Aif1-206
Aif1-201
Aif1-202
Aif1-204
Aif1-207
Ncr3-ps-201
Lst1-201
Ltb-201
Ltb-202
Ltb-203
Mir6973a-201
Tnf-201
Lta-201
Tnf-202
Mir6974-201
Nfkbil1-201
Gm16181-201
Nfkbil1-202
Atp6v1g2-201
Atp6v1g2-206
Atp6v1g2-203
Atp6v1g2-202
Atp6v1g2-205
Atp6v1g2-204
Ddx39b-205
Ddx39b-204
Ddx39b-206
Gm22589-201
Mir6975-201
Gm25128-201
SATB1 HiChIP
SATB1 binding sites
CTCF binding sites
H3K4me3 WT thymus ENCFF200ISF
H3K4me1 WT thymus ENCFF085AXD
H3K27ac WT DP cells GSM1504384+5
H3K27ac Rad21-cKO DP cells GSM1504386+7
ATACseq WT thymus
ATACseq Satb1-cKO thymus
RNAseq WT thymus
RNAseq Satb1-cKO thymus
Ltb
Tnf
Lta
B
chr11
99,000
99,025
99,050
99,075
99,100
99,125
99,150
99,175 Kb
99,200
Top2a-201
Top2a-206
Top2a-205
Top2a-204
Gm23451-201
Top2a-202
Igfbp4-201
Igfbp4-205
Igfbp4-206
Igfbp4-203
Igfbp4-202
Igfbp4-204
Tns4-201
Tns4-203
Tns4-202
Tns4-204
Ccr7-201
SATB1 HiChIP
SATB1 binding sites
CTCF binding sites
H3K4me3 WT thymus ENCFF200ISF
H3K4me1 WT thymus ENCFF085AXD
H3K27ac WT DP cells GSM1504384+5
H3K27ac Rad21-cKO DP cells GSM1504386+7
ATACseq WT thymus
ATACseq Satb1-cKO thymus
RNAseq WT thymus
RNAseq Satb1-cKO thymus
Ccr7
